# Human promoter directionality is determined by transcriptional initiation and the opposing activities of INTS11 and CDK9

**DOI:** 10.1101/2023.11.02.565285

**Authors:** Joshua D Eaton, Jessica Board, Lee Davidson, Chris Estell, Steven West

## Abstract

RNA polymerase II (RNAPII) transcription initiates bidirectionally at many human protein-coding genes. Sense transcription usually dominates and leads to messenger RNA production, whereas antisense transcription rapidly terminates. The basis for this directionality is not fully understood. Here, we show that sense transcriptional initiation is more efficient than in the antisense direction, which establishes initial directionality. After transcription begins, the opposing functions of Integrator (INTS11) and cyclin-dependent kinase 9 (CDK9) maintain directionality. Specifically, INTS11 terminates antisense transcription, whereas sense transcription is protected from INTS11-dependent attenuation by CDK9 activity. Strikingly, INTS11 attenuates transcription in both directions upon CDK9 inhibition, and the engineered recruitment of CDK9 desensitises transcription to INTS11. Therefore, the preferential initiation of sense transcription and the opposing activities of CDK9 and INTS11 explain mammalian promoter directionality.

## INTRODUCTION

In humans, most protein-coding genes initiate transcription bidirectionally. Usually, only sense transcription efficiently elongates and leads to mRNA synthesis. Antisense transcription terminates within a few kilobases (kb), and the short non-coding (nc)RNA is degraded^1^. Similar asymmetry frequently occurs in unicellular eukaryotes (e.g., budding yeast) and at some plant promoters^2,3^. Thus, promoter directionality is a broadly observed phenomenon. Interestingly, bidirectionality is the ground state of promoters and directionality is acquired over evolutionary time^4^. This is proposed to be through a combination of DNA sequences and proteins that favour the directional initiation and elongation of transcription.

The present explanation for the directionality of mammalian RNAPII promoters involves the arrangement of U1 snRNA binding sites and polyadenylation signals (PASs). U1 snRNA promotes RNAPII elongation through protein-coding genes by binding to RNA and preventing early termination^5–7^. RNA-bound U1 prevents early termination by inhibiting PASs and antagonising other attenuation mechanisms, which include the PP1 regulator, PNUTS, and the Restrictor complex^8,9^. In contrast, U1 binding sites are rarer in short antisense transcripts, which are often rich in PAS sequences that are proposed to promote transcriptional termination^10^. This model predicts that polyadenylation factors control a large fraction of antisense transcriptional termination.

The multi-subunit Integrator complex also regulates promoter-proximal transcription^11–15^. The Integrator complex comprises the backbone, arm/tail, phosphatase, and endonuclease modules^11^. Its endoribonuclease is INTS11, and its phosphatase activity is mediated by INTS6 and protein phosphatase 2A (PP2A). INTS11 endonuclease broadly affects promoter-proximal transcriptional attenuation whereas INTS phosphatase is proposed to regulate the escape of RNAPII into elongation^16,17^. INTS6/PP2A phosphatase functionally antagonises CDK9, which is vital for RNAPII promoter escape and productive elongation^18^. By this model, CDK9 activity promotes elongation across protein-coding genes and INTS6/PP2A opposes it.

We tested the prediction that PAS factors control transcription directionality by terminating antisense transcription, but this is not usually the case. Instead, we find that promoter directionality is often conferred by preferential initiation in the sense direction and is thereafter maintained by INTS11 and CDK9. The termination of antisense transcription is constitutively INTS11-dependent, whereas sense transcription is hypersensitive to INTS11 only when CDK9 activity is simultaneously inhibited. We hypothesise that CDK9 activity protects sense transcription from attenuation by INTS11 and that reduced CDK9 activity in the antisense direction exposes RNAPII to INTS11-dependent termination.

## RESULTS

### Polyadenylation factor depletion does not increase or extend antisense transcription

The current explanation of mammalian promoter directionality invokes early PAS-dependent termination of antisense transcription^10^. This is partly based on the direct detection of polyadenylated antisense transcripts, which provides evidence that some of their transcriptional termination is PAS-dependent. However, there are non-polyadenylated antisense RNAs that might be attenuated in other ways^19^. To test the contribution of PAS-dependent termination toward antisense transcriptional termination and promoter directionality we tagged *RBBP6* with the dTAG degron. Because RBBP6 is required to activate the PAS cleavage machinery^20,21^, its depletion should inhibit any PAS-dependent transcriptional termination. Three homozygous *dTAG-RBBP6* clones were isolated (Supplemental Figure 1A) and tagged RBBP6 was efficiently depleted after exposure to the dTAGv-1 degrader (Figure 1A). To test the contribution of RBBP6 to nascent transcription, we used POINT (Polymerase Intact Nascent Transcript)-seq^22^, which maps full-length RNA extracted from immunoprecipitated RNAPII.

**FIGURE 1:**
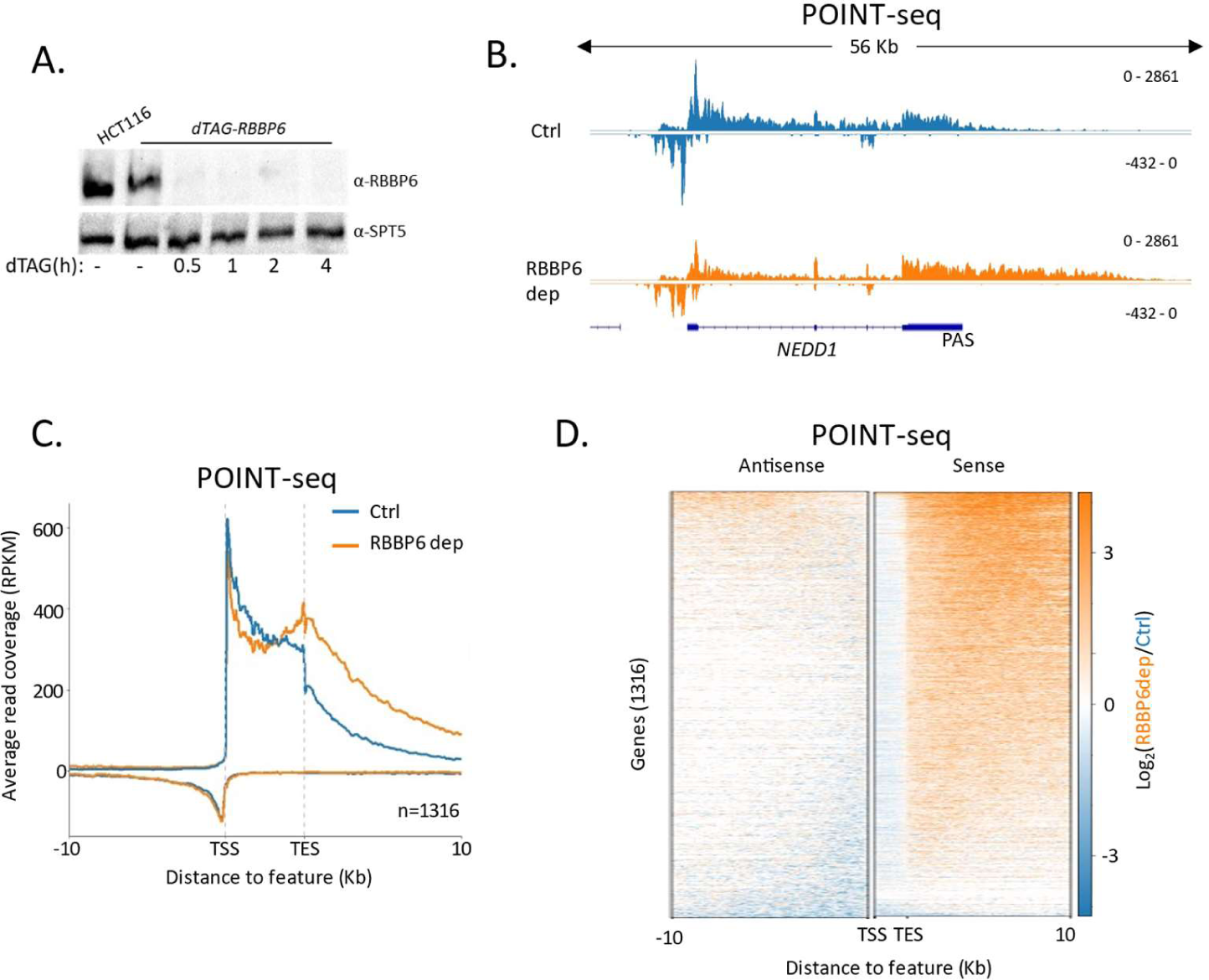
RBBP6 loss disrupts PAS-dependent termination of sense transcription. **A.** Western blot demonstrating the depletion of dTAG-RBBP6 over a time course of dTAGv-1 addition. SPT5 serves as a loading control. **B**. Genome browser track of *NEDD1* in POINT-seq data from *dTAG-RBBP6* cells treated or not (2hr) with dTAGv-1. RBBP6 depletion induces a termination defect in the protein-coding direction (downstream of the indicated PAS) but not the upstream antisense direction. The y-axis shows Reads Per Kilobase per Million mapped reads (RPKM). **C**. Metaplot of POINT-seq data from *dTAG-RBBP6* cells treated (RBBP6 dep) or not (Ctrl) (2hr) with dTAGv-1. This shows 1316 protein-coding genes selected as separated from any expressed transcription unit by ≥10kb. Signals above and below the x-axis are sense and antisense reads, respectively. The y-axis scale is RPKM. TSS=transcription start site; TES=transcription end site (this marks the PAS position). This is an average of two biological replicates. **D.** Heatmap representation of the data in C, which displays signal as a log2 fold change (log2FC) in RBBP6 depleted versus un-depleted conditions. This is an average of two biological replicates.

Figure 1B shows POINT-seq coverage over *NEDD1* including the upstream antisense transcript. RBBP6 loss causes a termination defect beyond the PAS shown by the extended POINT-seq signal beyond the *NEDD1* gene. However, RBBP6 does not affect the termination of upstream antisense RNA which is not extended when RBBP6 is depleted. Because RBBP6 loss induces strong termination defects beyond the PAS, we generalised its effects using genes separated from their neighbours by ≥10kb. Metaplot and heatmap analyses across these 1316 genes confirm that RBBP6 loss causes a general termination defect downstream of protein-coding genes but has little impact on antisense transcription (Figures 1C and D). Although many antisense transcripts contain multiple AAUAAA sequences, most still terminate RBBP6-independently (Supplementary Figures 1B and C). Similarly, we recently showed that the PAS-dependent 5’→3’ exonucleolytic torpedo terminator, XRN2, does not affect antisense transcriptional termination^9^. Therefore, although some antisense transcripts are polyadenylated, most antisense transcription can terminate using PAS-independent mechanisms. Consequently, PAS-dependent termination cannot fully explain the promoter-proximal attenuation of antisense transcription.

### Integrator depletion increases and extends antisense transcription

Our data argue that PAS-independent termination mechanisms control a large fraction of antisense transcription. A major PAS-independent termination pathway is driven by the Integrator complex, which was first identified as the 3’ end processing complex for snRNAs^23^.

Several reports show that Integrator terminates transcription from most promoters, including those that initiate antisense transcription, so it might affect promoter directionality^11,16,17^. To analyse this, we tagged its endonucleolytic subunit (INTS11) with a dTAG degron^24^, which enables rapid depletion of its catalytic activity (Figure 2A). We then performed POINT-seq on *INTS11-dTAG* cells depleted or not of INTS11 to assay the global impact of INTS11 on RNAPII transcription. Integrator is established to control the transcription of snRNAs, and this was detected in our POINT-seq, which demonstrates a clear extension of read density at *RNU5A-1* and *RNU5B-1* loci (Figure 2B).

**FIGURE 2:**
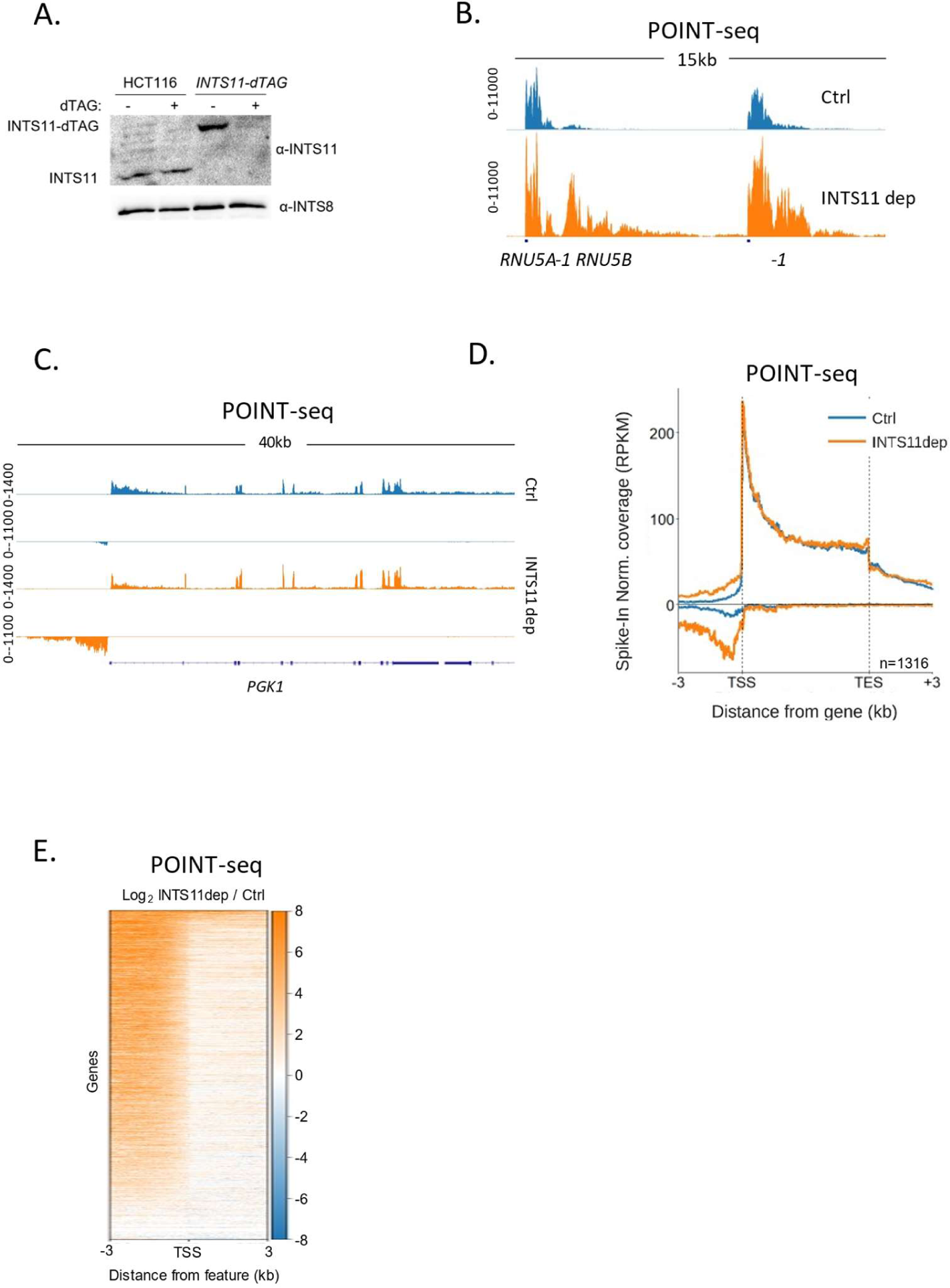
INTS11 loss disrupts the termination of antisense transcription. **A.** Western blot demonstrating homozygous tagging of *INTS11* with dTAG and the depletion of INTS11-dTAG after 1.5hr treatment with dTAGv-1. INTS8 serves as a loading control. **B**. Genome browser track showing POINT-seq signal over *RNU5A-1* and *RNU5B-1* in *INTS11-dTAG* cells treated or not (1.5hr) with dTAGv-1. Note, that INTS11 depletion induces a termination defect in each case. Y-axis shows RPKM following spike-in normalisation. **C**. Genome browser track showing POINT-seq signal over *PGK1* and its upstream antisense region in *INTS11-dTAG* cells treated or not (1.5hr) with dTAGv-1. Y-axis shows RPKM following spike-in normalisation. **D**. Metaplot of POINT-seq data from *INTS11-dTAG* cells treated or not (1.5hr) with dTAGv-1. This shows the same 1316 genes used in Figure 1C. Signals above and below the x-axis are sense and antisense reads, respectively. Y-axis shows RPKM following spike-in normalisation. This is an average of three biological replicates. **E.** Heatmap representation of the data in D, which displays signal as a log2 fold change (log2FC) in INTS11 depleted versus undepleted conditions over a region 3kb upstream and downstream of annotated TSSs. This is an average of three biological replicates.

As exemplified by *PGK1* (Figure 2C), INTS11 loss does not affect the termination of protein-coding transcription but causes a strong upregulation of antisense transcription. Meta-analysis of the same gene set used for RBBP6 POINT-seq shows the generality of antisense transcriptional attenuation via INTS11 (Figure 2D). A heatmap analysis of transcription 3kb upstream and downstream of the same transcription start sites (TSSs) demonstrates the dominant impact of INTS11 on antisense vs. sense transcription at most of these promoters (Figure 2E). Although the transcription over protein-coding genes is less affected by INTS11 loss, an increase in promoter-proximal transcription is apparent in some cases (Supplemental Figure 2A). The most strongly affected genes are lowly expressed, which is consistent with recent findings (Supplemental Figure 2B)^12–17^. Overall, INTS11 frequently attenuates antisense transcription whereas a smaller fraction of sense transcription is affected. RNAPII.

The hypersensitivity of antisense transcripts to INTS11 might be due to the makeup of antisense RNAPII or some other promoter feature. To interrogate this further, we analysed two other promoter classes: those where protein-coding transcripts are initiated in both directions and those that initiate the bidirectional transcription of unstable enhancer (e)RNAs. In the former case, both transcripts are extended and stable like most sense transcripts at directional promoters. In the latter case, both are short and unstable like most antisense transcripts at directional promoters. When both directions are protein-coding, INTS11 depletion causes modest reductions in transcription (Supplemental Figure 2C). Conversely, eRNA transcription is bidirectionally upregulated upon INTS11 elimination like the antisense transcripts in (Supplemental Figure 2D). We conclude that short ncRNAs are more strongly affected by INTS11 than protein-coding transcripts. At directional promoters this results in the attenuation of antisense transcription.

### Transcription initiates more efficiently in the sense direction

The increased antisense POINT-seq signal following INTS11 loss is consistent with defects in transcriptional termination. However, it could result from increased transcriptional initiation. We were also interested in whether preferential sense initiation could contribute to mammalian promoter directionality as was proposed in budding yeast^4^. To precisely resolve directional aspects of initiation and assay any impact of INTS11, we devised a variant of POINT-seq called short (s)POINT. Briefly, sPOINT follows the POINT-seq protocol, but library preparation employs the selective amplification of 5’ capped RNAs <150nts (Figure 3A, Supplemental Figure 3A, Experimental Procedures). The sPOINT signal over *PGK1* exemplifies this and demonstrates a very restricted signal close to the TSS (Figure 3B). Figure 3C compares the meta profile of POINT- and sPOINT-seq on 684 well-expressed and well-spaced (≥10kb from neighbours) genes to highlight the full read coverage obtained by POINT-seq and the tightly restricted, TSS-proximal, sPOINT signal. Because most capped RNAPII-associated RNA <150nts are promoter-proximal, sPOINT effectively assays the promoter-proximal RNAPII pause.

**FIGURE 3:**
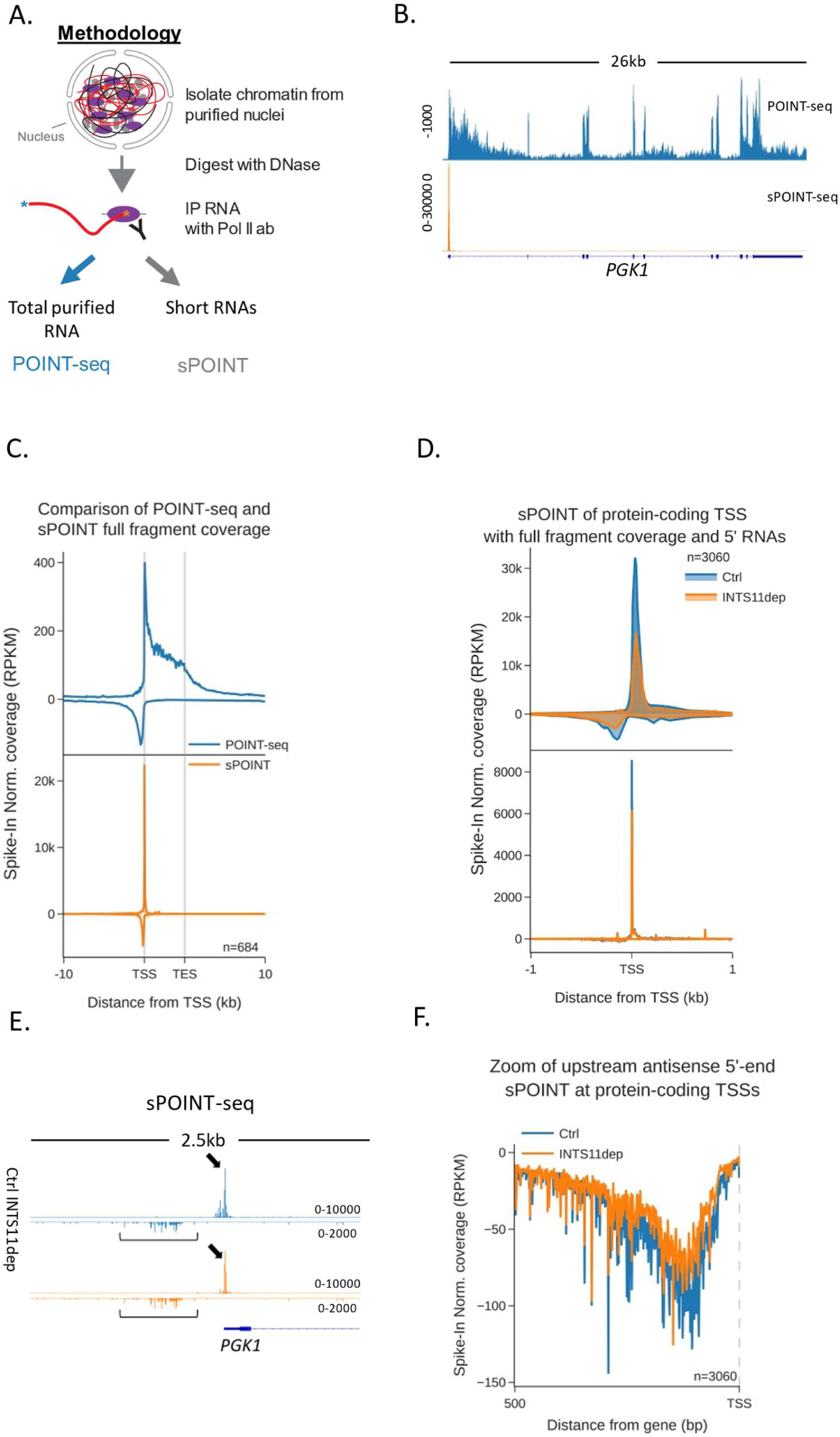
Sense direction transcription initiation is more efficient and focused. **A.** Schematic of sPOINT-seq protocol. The POINT-seq protocol is followed, in which chromatin is isolated and engaged RNAPII is immunoprecipitated. Short transcripts are preferentially amplified during library preparation (see Experimental Procedures for full details). **B.** Comparison of POINT-(top trace) and sPOINT-seq (lower trace) on *PGK1*. Y-axis units are RPKM. **C**. Metaplot comparison of POINT-(top plot) and sPOINT-seq (lower plot) profiles across the 684 highest expressed protein-coding that are separated from expressed transcription units by ≥10kb (top ∼50%). Signals above and below the x-axis are sense and antisense reads, respectively. Y-axis shows RPKM following spike-in normalisation. **D.** Top metaplot shows full read coverage for sPOINT-seq performed in *INTS11-dTAG* cells treated or not (1.5hr) with dTAGv-1 at the promoters of the top expressed 20% of protein-coding genes. The lower metaplot is the same data but only the 5’ end of each read is plotted. The y-axis signals are RPKM following spike-in normalisation. Two biological replicates of sPOINT were performed. **E**. Genome browser track of *PGK1* promoter region in sPOINT-seq. This showcases the focused sense TSS (black arrows) and the dispersed antisense reads (brackets). Note the higher y-axis scale (RPKM) for sense vs. antisense. **F**. Metaplot zoom of the antisense TSS signals deriving from the lower plot in D. This makes clear the dispersed sites of initiation. The Y-axis scale is RPKM following spike-in normalisation. Two biological replicates of sPOINT were performed.

The sPOINT-seq metaplot in Figure 3B shows a higher signal in the sense direction, suggesting that more efficient transcriptional initiation establishes promoter directionality. To assay this further and test the impact of INTS11, we performed sPOINT-seq in *INTS11-dTAG* cells treated or not with dTAGv-1 and plotted the coverage over the promoters of well-expressed protein-coding genes (3060 promoters, Figure 3D). As sPOINT-seq maps the 5’ and 3’ ends of these reads, precise TSSs are mapped at single-nucleotide resolution and plotted in the lower meta profile in Figure 3D. This analysis once again shows a higher sPOINT signal in the sense direction. In addition, the lower TSS mapped plot also shows that sense transcription initiated in a more focused manner compared to in the antisense direction. Figure 3E (*PGK1*) and supplemental figure 3B (*ACTB*) exemplify these features on individual genes, and the metaplot in Figure 3F highlights the dispersed nature of antisense TSSs. Quantitation of the TSS-derived sense vs. antisense signal confirms a higher read count in the sense direction in untreated cells (Supplemental Figure 3C). The most noticeable (although still modest) effect of INTS11 loss on sPOINT-seq profiles is a mild signal reduction. This shows that INTS11 depletion does not generally enhance transcriptional initiation in either direction. This slight reduction in sPOINT-seq signal could result from less transcriptional initiation or less efficient pausing if, for example, INTS11 loss allows more RNAPII to escape the promoter. Although a lower resolution technique, RNAPII ChIP-seq confirmed these promoter characteristics and the mild impact of INTS11 (Supplemental Figures 3D and E). Overall, these data show that directionality might be established by more efficient initiation in the sense direction.

### CDK9 inhibition sensitises sense transcription to INTS11

After transcription initiates, further control of directionality is evident because sense transcription goes further than antisense transcription. If INTS11 regulates this, an opposing force is required to promote elongation. INTS11 affects transcription very early, occupies promoters, and becomes less active as RNAPII moves into elongation^16–18,25,26^. Therefore, if INTS11 is counteracted to allow sense transcription, any responsible mechanism needs to act early. One of the first transcriptional checkpoints involves the phosphorylation of RNAPII and other factors by CDK9, which releases promoter-proximally paused RNAPII into elongation^27^. During this process, the Integrator-associated phosphatase, PP2A, antagonizes CDK9 and presumably regulates the sensitivity of RNAPII to INTS11^18^. Because Integrator phosphatase has little effect on antisense transcription^16^, we hypothesised that INTS11 sensitivity and CDK9 activity are inversely correlated to maintain directionality after initiation.

Our hypothesis predicts that sense transcription will be attenuated via INTS11 when CDK9 is inactive. To test this genome-wide, we depleted INTS11 from *INTS11-dTAG* cells in the presence or absence of a specific CDK9 inhibitor (NVP-2^28^) and performed POINT-seq. As exemplified by *TARDBP*, the depletion of INTS11 alone caused an antisense transcriptional termination defect with a milder impact on the protein-coding sense direction (Figure 4A). As expected, NVP-2 treatment reduced transcription over the protein-coding gene body. Antisense transcription also displays some CDK9 sensitivity. Importantly, in NVP-2-treated cells, INTS11 loss increased transcription in both directions. This contrasts with the dominant antisense effect deriving from just depleting INTS11 (refer to Figure 2). As such, CDK9 activity prevents sense transcription from being attenuated by INTS11. This is a genome-wide trend as shown in the metaplots in Figures 4B and C. A heatmap of the effect of INTS11 loss after CDK9 inhibition shows the bidirectional upregulation of transcription at the same promoters assayed in Figure 2 (Figure 4D). Although this experiment employed longer INTS11 depletion than that in Figure 2 (2.5hr vs. 1.5hr to allow concurrent CDK9 inhibition), its antisense effects remain general, and the affected protein-coding genes strongly overlapped (Supplemental Figure 4A-C). Lastly, when CDK9 and INTS11 are both compromised, the POINT-seq signal remains higher in the sense vs. antisense direction (black line, Figure 4C). This is consistent with our sPOINT-based finding that initiation of sense transcription is generally more efficient.

**FIGURE 4:**
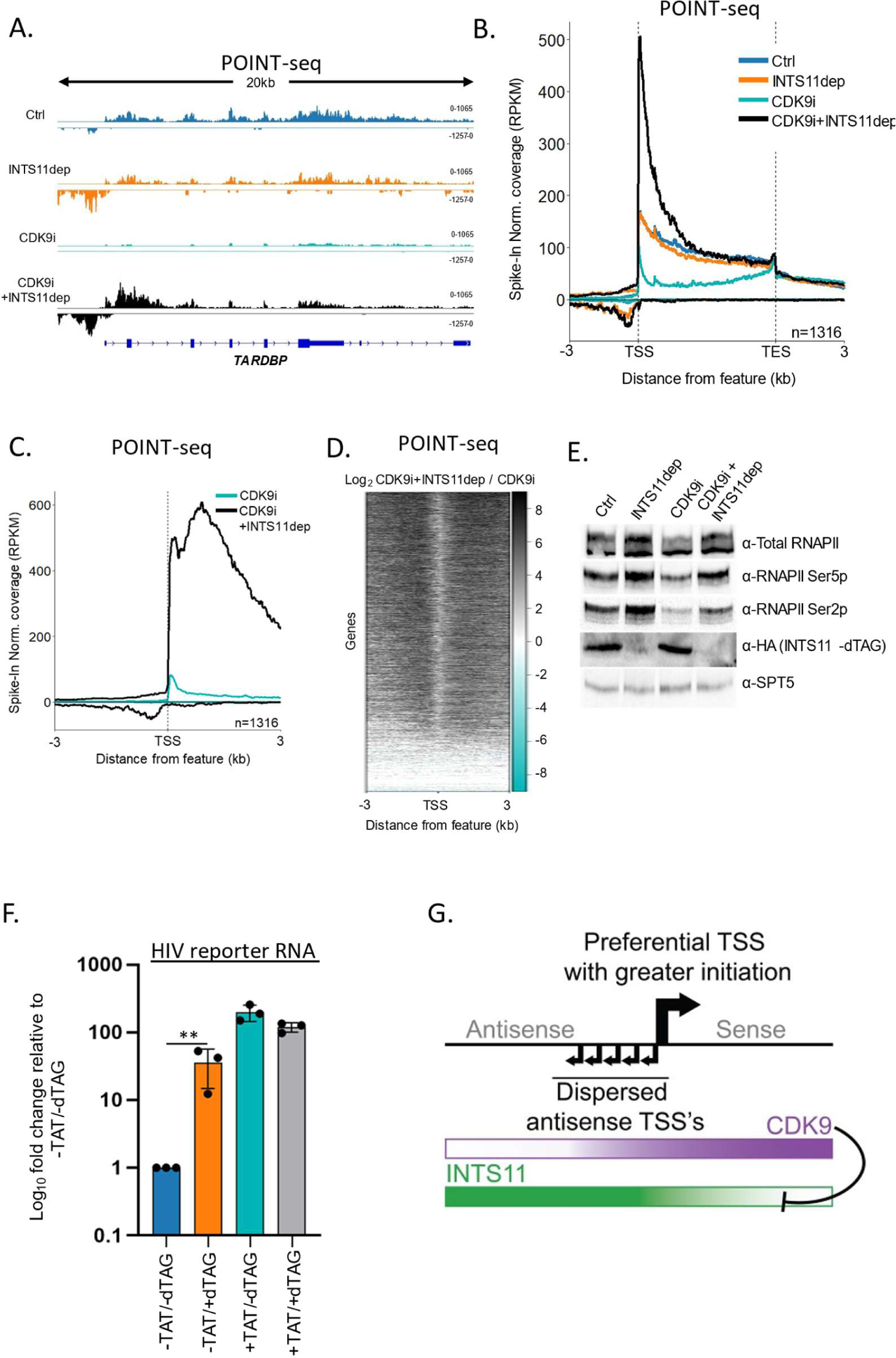
Some promoter directionality is lost when CDK9 is inhibited. **A**. Genome browser track of *TARDBP* in POINT-seq data derived from *INTS11-dTAG* cells either untreated, dTAG treated, NVP-2 treated or dTAG and NVP-2 treated (2.5 hr). Signals above and below the Y-axis are sense and antisense reads, respectively. The Y-axis scale shows RPKM following spike-in normalisation. **B**. Metaplot of POINT-seq in *INTS11-dTAG* cells depleted or not of INTS11 and treated or not with NVP-2 (2.5 hr). This uses the same gene set as Figure 1C. The regions 3kb upstream and downstream of genes are included. Y-axis units are RPKM following spike-in normalisation. This is an average of two biological replicates. **C**. Metaplot of the same CDK9i + and -dTAG data shown in B but zoomed into the region 3kb upstream and downstream of the TSS. This is an average of three biological replicates. **D.** Heatmap representation of the data in C, which displays signal as a log2 fold change (log2FC) in INTS11 depleted versus un-depleted conditions covering a region 3kb upstream and downstream of the TSS. This is an average of three biological replicates. **E.** Western blot for total RNAPII and RNAPII phosphorylated on Ser2/5 (Ser2/5p) in *INTS11-dTAG* cells treated or not with dTAGv-1 and/or NVP-2 (as indicated). dTAG-mediated INTS11 depletion is shown in the anti-HA blot and SPT5 us used as a loading control. **F.** qRT-PCR analysis of *INTS11-dTAG* cells transfected with the HIV reporter construct with or without TAT then depleted or not of INTS11. Quantitation shows signals relative to those obtained in the presence of INTS11 and the absence of TAT after normalising to MALAT1 RNA. n=3. Error bars show standard deviation. ** denoted p=0.01. Note that INTS11 depletion was performed concurrently with transection (14hr in total). **G.** Model for promoter directionality depicting higher levels of focused transcriptional initiation in the sense direction together with opposing gradients of CDK9 and INTS11 activity that peak in sense and antisense directions, respectively.

Following CDK9 inhibition and INTS11 loss, the largest recovery of protein-coding transcription is often over the 5’ end of genes. This presumably reflects poor elongation without CDK9 activity even when INTS11 is absent. RNAPII elongation is associated with phosphorylation of the C-terminal domain (CTD) of its largest subunit, RPB1, and most frequently occurs on Serine 5 or 2 (Ser5/2p) of its heptad repeat^29,30^. We assayed the effects of CDK9 inhibition and INTS11 depletion on these two modifications by western blotting (Figure 4E). As previously shown^28^, CDK9 inhibition by NVP-2 suppresses Ser2p. However, both Ser5p and Ser2p are enhanced by INTS11 loss whether CDK9 is active or not. Thus, unattenuated transcription after INTS11 depletion is associated with some CTD phosphorylation. Where NVP-2 is employed, kinases redundant with CDK9 presumably phosphorylate RNAPII.

We confirmed the CDK9-mediated suppression of INTS11 on four selected protein-coding genes using an alternative inhibitor, 5, 6-dichloro-1-β-D-ribofuranosylbenzimidazole (DRB) (Supplemental Figure 4D). Interestingly, inhibition of CDK7 with THZ2 sensitised the same selected protein-coding transcripts to INTS11 attenuation (Supplemental Figure 4E). This may be because CDK7 is required to activate CDK9^31^, or because Integrator additionally targets RNAPII when CDK7 is inactive. This result reinforces the idea that the successful execution of early transcriptional checkpoints may overcome Integrator-mediated attenuation.

If CDK9 counteracts INTS11, its recruitment should prevent transcriptional attenuation by Integrator. To assay this, we employed a plasmid where transcription is driven by the human immunodeficiency virus (HIV) promoter. Transcription from the HIV promoter results in synthesis of the trans-activating response (TAR) element and promoter-proximal RNAPII pausing. Pause release requires the trans-activator of transcription (TAT), which promotes RNAPII elongation by recruiting CDK9^32^. INTS11 suppresses transcription from the HIV promoter when TAT is absent^33^. To assay whether CDK9 affects this process, *INTS11-dTAG* cells were transfected with the HIV reporter with or without TAT before treatment or not with dTAGv-1. Transcription from the reporter was then analysed by qRT-PCR (Figure 4F). INTS11 loss induces HIV transcription in the absence of TAT. However, TAT strongly stimulates transcription (∼200 fold) and desensitises it to INTS11 loss (slightly less reporter RNA was recovered). Therefore, TAT-mediated CDK9 recruitment alleviates INTS11-dependent attenuation of transcription. In line with endogenous protein-coding genes, CDK9 inhibition sensitises TAT-activated transcription to INTS11 (Supplemental Figure 4F). Overall, our data strongly suggest that CDK9 activity counteracts transcriptional attenuation by INTS11.

## DISCUSSION

Most studies on mammalian promoter directionality have focused on understanding how antisense transcription terminates and how RNAPII elongation is enabled in the sense direction. However, we show that preferential sense transcriptional initiation significantly contributes toward the directionality of many mammalian promoters. More analyses are required to elucidate the mechanism, but we hypothesise that a unidirectional arrangement of at least some core promoter elements favours transcription in the protein-coding direction. Consistently, the evolution of promoter elements explains how a bidirectional promoter “ground state” acquired directionality in yeast^4^. Consistent with our finding, focused transcription initiation is associated with highly expressed mammalian genes and promoters with clearly defined elements (e.g. TATA)^34^. Thus, the lower level and dispersed nature of antisense initiation may reflect the suboptimal orientation of promoter elements or the opportunistic transcription of open chromatin in promoter regions.

We hypothesise that INTS11 controls transcriptional attenuation via its endonuclease activity. An inactive mutant of INTS11 (E203Q) is widely used to interrogate its catalytic function^23^; however, we discovered that it poorly associates with other Integrator components compared to wild-type INTS11 (Supplemental Figure 4G). Therefore, when employed in cells, it may not isolate the effects of INTS11 activity from those requiring an intact Integrator complex. Similarly, catalytic mutations in the highly related PAS endonuclease, CPSF73, disrupt its association with other cleavage and polyadenylation components^35^. In light of this, it is formally possible that non-catalytic consequences of INTS11 loss explain the upregulation of antisense transcription.

How CDK9 activity opposes INTS11 is unresolved, but INTS11 and SPT5 are adjacent in the RNAPII: Integrator structure^36^. As SPT5 is a prominent substrate of CDK9^37–39^, its phosphorylation might evict INTS11 or prevent its association with the complex. Consistently, Integrator is enriched on promoters whereas phosphorylated SPT5 (SPT5p) is most prevalent during elongation over protein-coding gene bodies^25,38^. Furthermore, although antisense RNAPII occupancy is increased by INTS11 loss, published data suggest that this is not paralleled by increases in its Ser5p and Ser2p forms^16^. As these modifications require CDK7 and CDK9, this provides further support that these activities are less prevalent for antisense vs. sense transcription.

While INTS11 and CDK9 activities are the clearest on antisense and sense transcription, respectively, this is not binary. Some antisense transcription is NVP-2 sensitive, which indicates CDK9 activity on some RNAPII (Figure 4 and^40^). CDK9-sensitive antisense transcription could undergo INTS11-independent attenuation, which might account for the polyadenylated antisense RNA observed previously^10,41^. Consistently, some antisense transcription is targeted by the poly(A) exosome targeting connection, which operates on cleaved and polyadenylated transcripts^42,43^. Reciprocally, promoter-proximal transcription from lowly expressed protein-coding transcription is INTS11-sensitive, which could reflect low CDK9 activity at those loci. We and others recently described the Restrictor complex as restraining antisense transcription^9,44,45^. Unlike Integrator, which associates with NELF-bound RNAPII^36^, Restrictor is proximal to the distinct PAF-bound RNAPII^9^. Thus, various complexes may target different forms of RNAPII before and after CDK9 activity. Even so, our data suggests that Integrator influences a greater volume of antisense RNAPII than the cleavage and polyadenylation pathway.

In addition to CDK9, multiple studies demonstrate a role for U1 snRNA in promoting elongation through protein-coding genes^5,6,8,46^. U1 promotes RNAPII elongation and shields transcription from early PAS-dependent termination and the Restrictor complex^5,9^. The U1-mediated suppression of premature PAS-dependent termination inspired the original model of promoter directionality. As U1 sites are rarer in antisense RNAs, their termination was hypothesised to be driven by early PASs that would consequently be active. Our RBBP6 data (and other published data on PAS termination factors^9^) argues that a large fraction of antisense transcriptional termination is PAS-independent. Nevertheless, the lack of U1 sites may limit RNAPII elongation and render it sensitive to Integrator (or Restrictor). Accordingly, it will be interesting to test whether U1 regulates the sensitivity of transcription to Integrator.

In conclusion, we provide a new model for mammalian promoter directionality and the subsequent control of bidirectional transcription (Figure 4G). sPOINT demonstrates more efficient sense transcription initiation, which establishes directionality. Early events dictate the decision to elongate or attenuate transcription. In the sense direction, CDK9 activity prevents attenuation by Integrator to favour productive elongation, whereas antisense transcription is hypersensitive to INTS11 and terminates early. As INTS11 occupies most promoters and CDK9 activity is near-universally required to achieve protein-coding transcription, this model can generally explain how directionality is initiated and maintained. Our new sPOINT-seq approach will be valuable in elucidating additional aspects of promoter-proximal transcriptional regulation.

## ACKNOWLEDGEMENTS

We thank Hiroshi Kimura for the total RNAPII antibody used for POINT-seq. Our research is funded by a Wellcome Trust Investigator Award to SW (223106/Z/21/Z). This project used the University of Exeter Sequencing Service, and their equipment was funded by the Wellcome Trust (Multi-User Equipment Grant award number 218247/Z/19/Z).

## AUTHOR CONTRIBUTIONS

Conceptualization J.D.E, S.W; data curation J.D.E, J.B, L.D, C.E; formal analysis J.D.E., J.B., L.D; C.E and S.W; methodology, J.D.E. and S.W.; investigation, J.D.E., J.B., L.D; C.E and S.W.; supervision and funding acquisition, S.W.; writing and editing, J.D.E., J.B., L.D; C.E and S.W.

## EXPERIMENTAL PROCEDURES

### Sequencing data

Deposited at Gene Expression Omnibus under accession: GSE243266.

### Cloning

HIV reporter constructs were made by removing the CMV promoter, the entire β-globin sequence, and its PAS from a pcDNA5 FRT/TO plasmid containing the WT β-globin (βWT) gene^47^ and inserting an HIV promoter and downstream TAR element derived from βΔ5-7^48^. *INTS11* targeting constructs were modified from those we previously described to generate *INTS11-SMASh* cells^49^. The SMASh tag was removed and replaced with 2xHA dTAG derived from Addgene plasmid 91792^24^. Guide RNA expressing Cas9 plasmids to modify *INTS11* or *RBBP6* were made by inserting annealed oligonucleotides, containing the targeting sequence, into px330^50^ digested with BbsI.

### Cell culture and cell lines

HCT116 cells were maintained in DMEM supplemented with penicillin/streptomycin at 37°C, 5% CO_2_. *dTAG-RBBP6* cells were generated using the “CHoP in” protocol^51^. Briefly, a 24-well dish was transfected with 250ng of px330^50^ containing the *RBBP6*-targeting guide and 250ng of PCR product containing the dTAG degron preceded by a blasticidin or puromycin selection marker (derived from Addgene plasmid 91792 and 91793^24^). Jetprime (Polyplus) was used for transfection. Three days later, cells were passaged into media containing 10µg/ml Blasticidin/1µg/ml Puromycin and colonies were PCR screened ∼10 days later. *INTS11-dTAG* cells were generated by homology-directed repair. A 6-well dish of cells was transfected with 1µg px330 containing the *INTS11*-targeting guide (described in^49^) and 1µg each of the repair templates. Three days later, cells were passaged into media containing 30µg/ml Hygromycin and 800µg/ml G418. ∼10 days later, colonies were picked and screened by PCR. 1µM dTAGv-1 (Tocris) was used for 1-14hrs (see figure legends for timings used in each experiment); NVP-2 was used at 250nM for 2.5hr; DRB was used at 100µM for 2.5hr; THZ2 was used at 5µM for 2.5hr.

### POINT-seq and sPOINT-seq

For POINT-seq, we followed the protocol provided in^22^. The only modification was that we started with a confluent 10cm dish of cells and performed the immunoprecipitation with 6µg of anti-RNAPII. ∼2% cell volume of Drosophila S2 cells was included as a spike in control. Libraries were prepared using the NEBNext Ultra™ II Directional RNA Library Prep Kit for Illumina (New England Biolabs). sPOINT was performed in the same manner with the following difference: a confluent 15cm dish of cells was used and 10µg of anti-RNAPII. In sPOINT, the immunoprecipitated and DNAse treated RNA was treated with Terminator™ 5-Phosphate-Dependent Exonuclease (lucigen) to remove any uncapped transcripts, and libraries were prepared with the SMARTer® smRNA-Seq Kit for Illumina (Takara Bio) to selectively capture transcripts <150nts.

### ChIP-seq

For each experiment, 1×10cm dish of cells was used. Protein: DNA crosslinks were formed by adding Formaldehyde (1% v/v) to culture media for 10 mins then quenching with 125mM glycine. Cells were rinsed 2x with PBS, scraped off the dish, and pelleted in 10ml PBS at 500xg for 5 mins. We then employed the simple ChIP enzymatic kit (Cell Signalling Technologies) to fragment chromatin and purify RNAPII-bound DNA. We followed the kit protocol except for conjugating 5µg of anti-total RNAPII to sheep anti-mouse dynabeads (Life Technologies). Sequencing libraries were generated using the NEBNext® Ultra™ II DNA Library Prep Kit for Illumina.

### Total/chromatin-associated RNA isolation and qRT-PCR

For total RNA, a 24-well dish of cells was transfected with 100ng HIV reporter plasmid and, where co-transfected, 50ng TAT plasmid^52^ using Jetprime (Polyplus) following the manufacturer’s protocol. Media was refreshed 5 hours post-transfection and dTAGv-1 was added where appropriate. The next day, total RNA was isolated using Trizol (Thermo Fisher) following the manufacturer’s protocol. For chromatin-associated RNA (for Supplemental Figure 4F), a 6 well dish of cells was transfected with 300ng HIV reporter plasmid and 150ng TAT. Pelleted cells were resuspended in 800µl hypotonic lysis buffer (HLB: 10 mM Tris-HCl (pH 7.5), 10 mM NaCl and 2.5 mM MgCl_2_, 0.5% NP40) and underlayered with 200µl HLB +10% sucrose. Nuclei were isolated by centrifugation for 5 mins at 500xg. These were resuspended in 100µl NUN1 (20 mM Tris-HCl (pH 7.9), 75 mM NaCl, 0.5 mM EDTA, and 50% Glycerol). After addition of 1ml of NUN2 (20 mM HEPES-KOH (pH 7.6), 300 mM NaCl, 0.2 mM EDTA, 7.5 mM MgCl_2_, 1% NP-40, 1 M Urea) chromatin was isolated by 10 min incubation on ice followed by 10 min centrifugation at 13000rpm. RNA was isolated from the chromatin pellet using Trizol following the manufacturers’ instructions. For qRT-PCR of total and chromatin-associated RNA, RNA was DNase treated then 1µg was reverse transcribed using Protoscript II reverse transcriptase (New England Biolabs). qPCR was performed using LUNA SYBR green reagent (New England Biolabs) on a Qiagen Rotorgene instrument. Quantitative analysis used the ΔCT method.

### Co-immunoprecipitation

For each transfection, a semi-confluent 10cm dish of cells was transfected with 5µg of plasmid expressing flag tagged wild type or E203Q INTS11. Immunoprecipitation was based on the ELCAP protocol^53^. The following day, cells were washed 2x in PBS, scrapped into 10ml PBS and spun down for 5 mins at 500xg. Pelleted cells were resuspended in 800µl hypotonic lysis buffer low NP40 (HLB: 10 mM Tris-HCl (pH 7.5), 10 mM NaCl and 2.5 mM MgCl_2_, 0.1% NP40) and underlayered with 200µl HLB low NP40 +10% sucrose. Nuclei were resuspended in 1ml Chromatin digestion buffer (20 mM HEPES pH 7.9, 1.5 mM MgCl2, 10% (v/v) glycerol, 150 mM NaCl, 0.1% (v/v) NP-40 and 250 U/mL Benzonase) and incubated for 1hr at 4°C. Debris was pelleted at 13000rpm for 10 mins and supernatant was incubated with 20µl of anti-flag magnetic beads (Sigma) for 2hrs at 4°C. Beads were washed 6x with ice-cold chromatin digestion buffer and samples were eluted by boiling in protein loading buffer (100 mM Tris-Cl (pH 6.8) 4% (w/v) sodium dodecyl sulfate 0.2% (w/v) bromophenol blue 20% (v/v) glycerol) before polyacrylamide separation and western blotting.

### qRT-PCR primers and gRNA target sites

See supplemental table 1.

## BIOINFORMATICS

### POINT-Seq alignment and visualisation

A metagene list of genes with no overlapping regions within 10kb of any other expressed transcription unit (1316). Adapters were removed from raw reads using Trim Galore! and mapped to GRCh38 using HISAT2 using default parameters. Biological replicates were normalised and merged using SAMtools merge. Split strand metagene plots were produced using Drosophila spike-in normalised sense and antisense (scaled to -1) bigwig coverage files separately with further graphical processing performed in R. For heat maps, computematrix (DeepTools) was used to generate score files from the normalised bigwig files using the 10kb non-overlapping gene list. A log2 ratio (depletion/control) was applied to identify changes in reads. Plots were redrawn in R; parameters used for each heat map are detailed in figure legends.

### sPOINT

For sPOINT TSS metaplots showing full-length and 5’ derived coverage, gene lists were determined by selecting principal protein-coding transcript isoforms from gencode v42 human annotation – specifically, those containing both “appris_principal_1” and “Ensembl_canonical” labels (15301 in total). The top 20% expressed (based on -dTAG) were used to generate meta profiles (3060 TSSs in total). The 684 genes used to exemplify the difference between POINT and sPOINT coverage are the top 50% of expressed genes (based on POINT-seq in *INTS11-dTAG)* cells, which are separated from neighbouring transcription usints by at least 10kb.

### Antisense PAS analysis

Bed files from Figure 1C were edited to map the antisense transcript from each gene by fitting each gene’s start and end point to 3kb upstream of their TSS. Sequences underlying these regions were obtained via BEDTools getFASTA (hg38) and consensus PAS (AATAAA) enumerated in R. PAS frequency per transcript was plotted using ggplot2. Heatmaps for transcripts containing > 1 or no PAS motifs were plotted with DeepTools.

### ChIP-seq alignment and plotting

Adapters were removed from raw reads using Trim Galore! and mapped to GRCh38 using HISAT2 with default parameters. Reads were also mapped to Dm6 to identify spike-in signal. Reads with MAPQ score of ≤30 were removed with SAMtools^54^. Peaks were called using MACS2 in paired-end mode^55^. Metaplots were created from S2 scaled, merged replicate -log10 q-value bigwig coverage using DeepTools with further processing performed within R. For Supplementary Figure 3E, plots used the same gene list described above for sPOINT promoter analyses (see sPOINT section above).

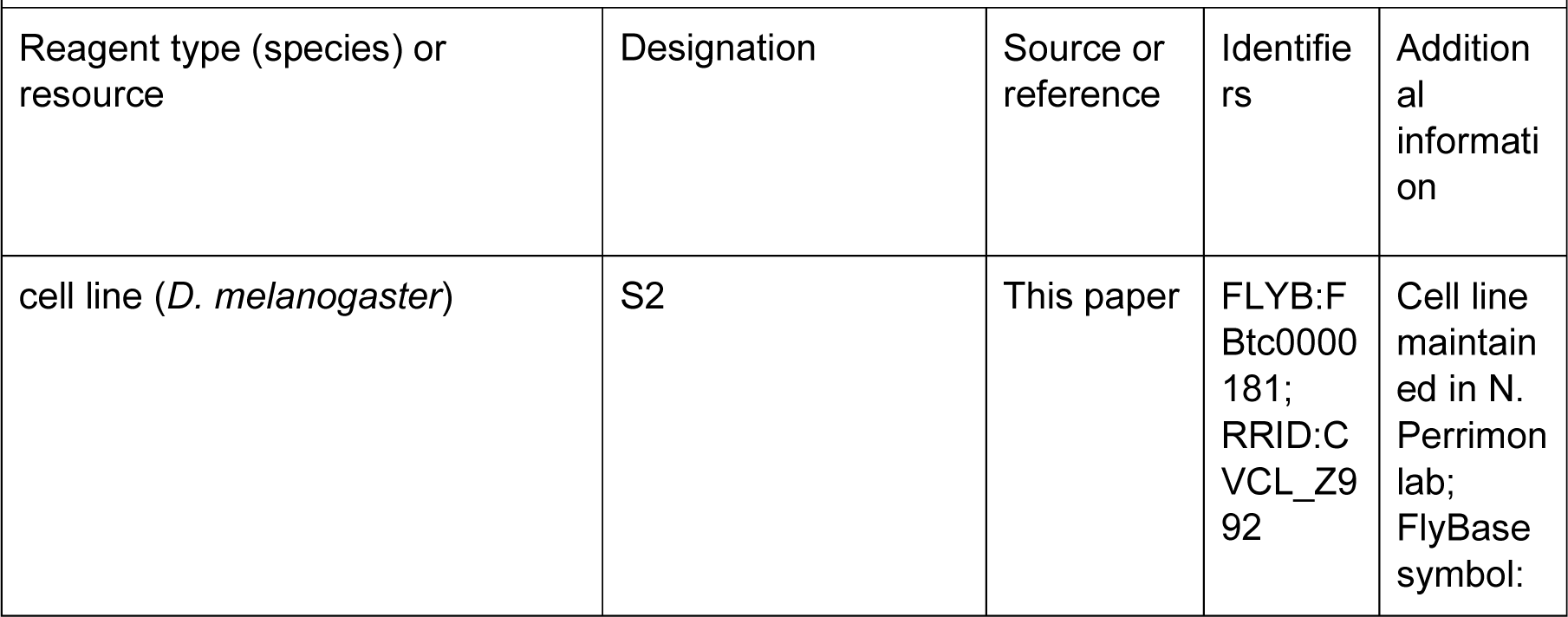
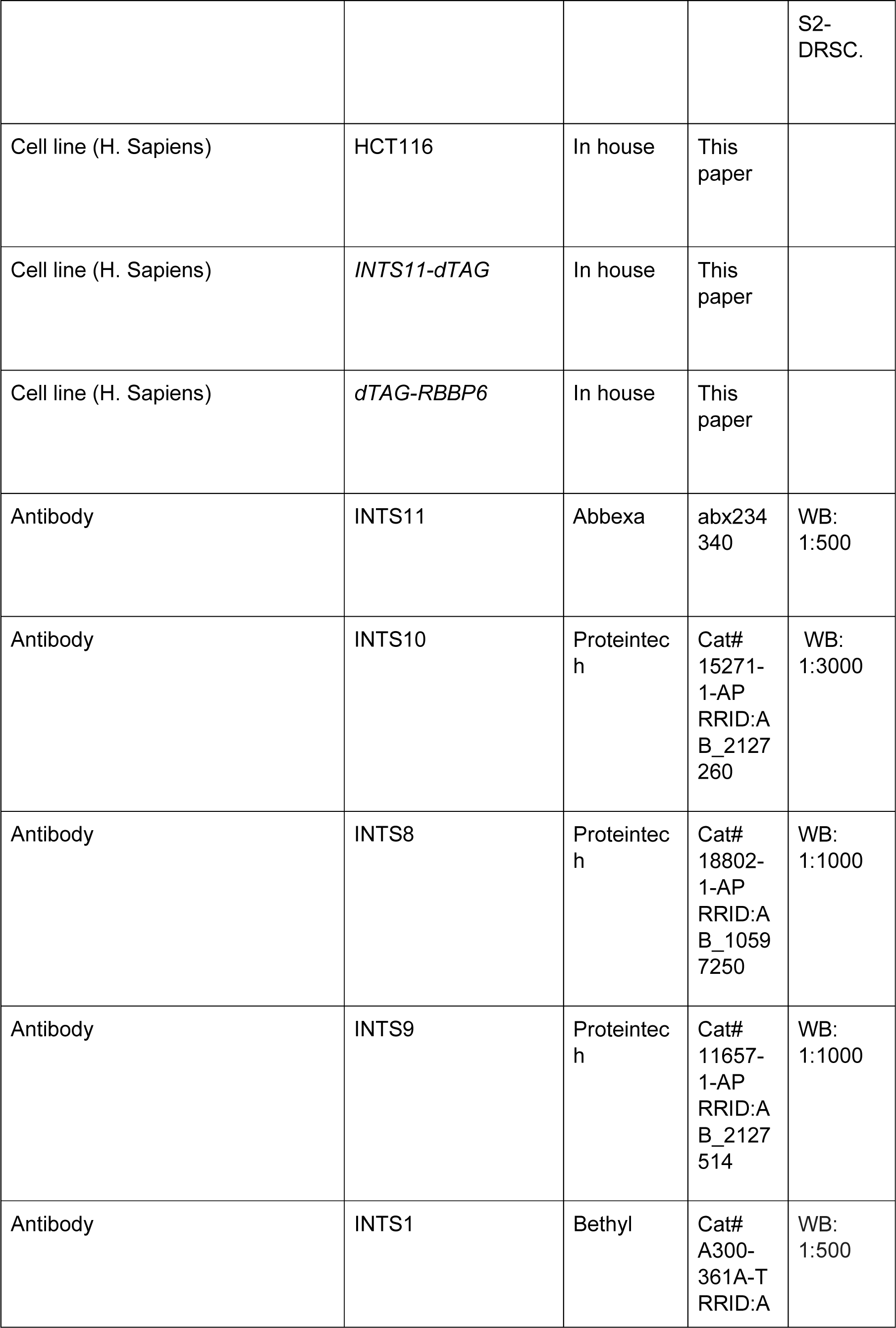
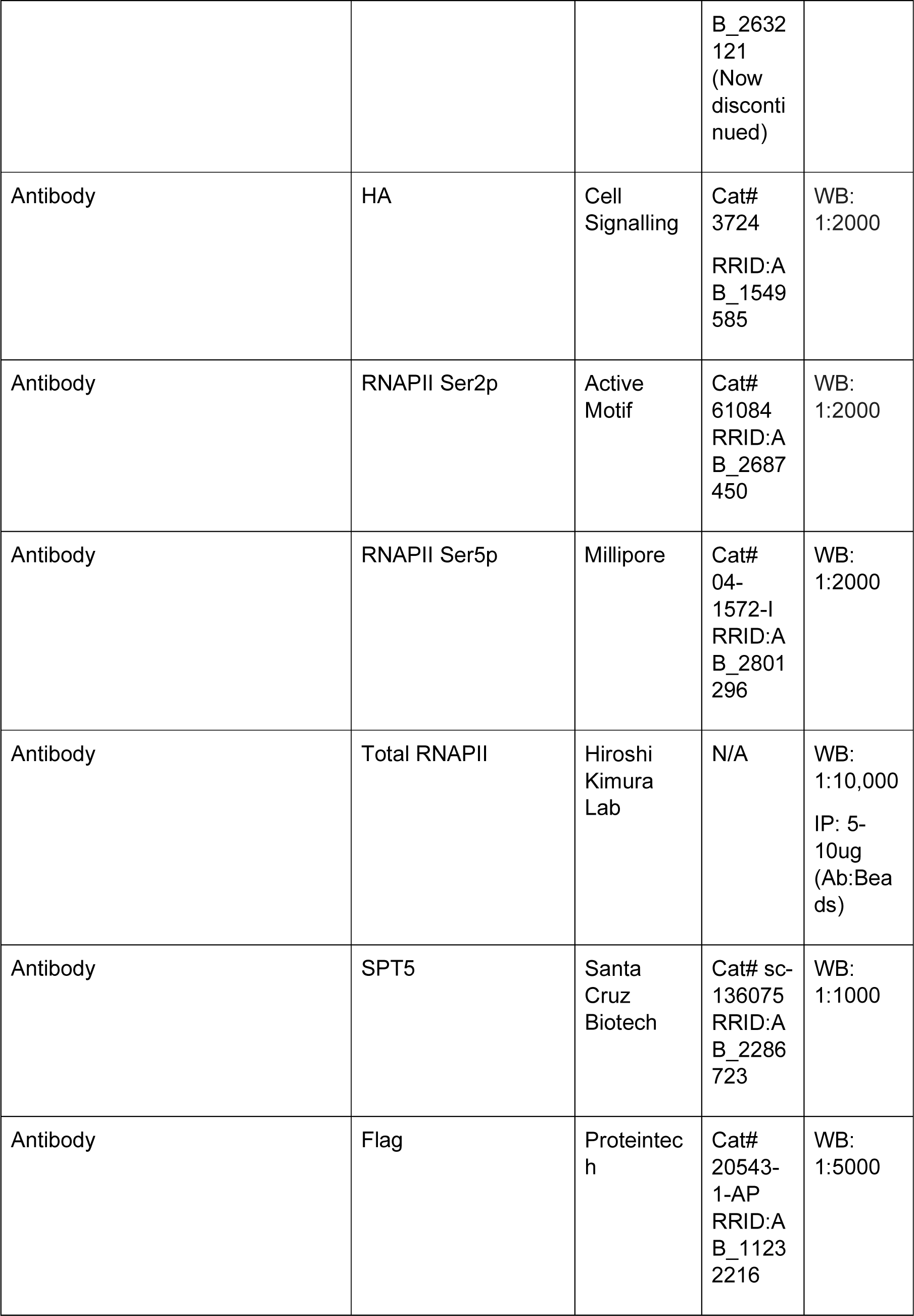
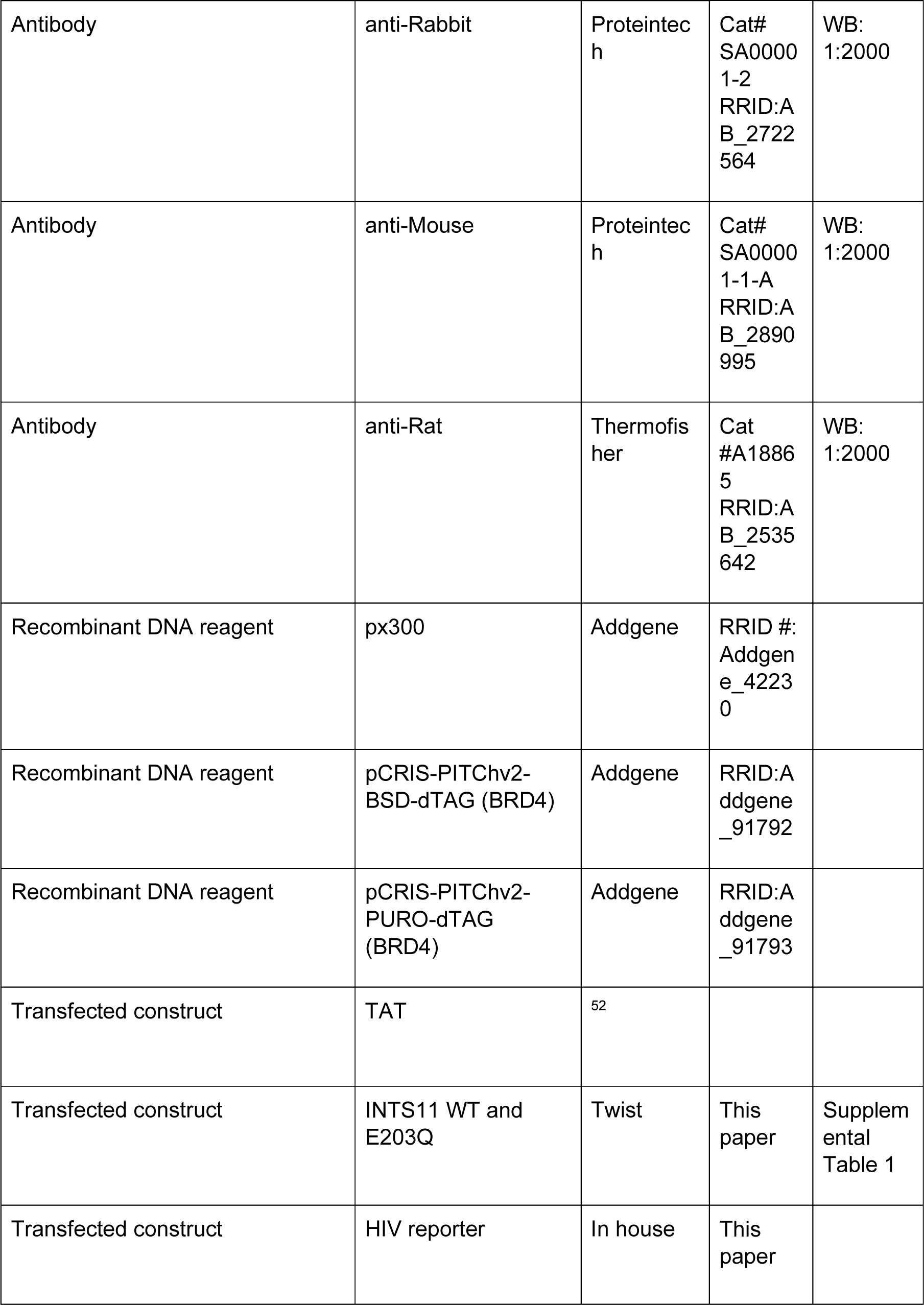
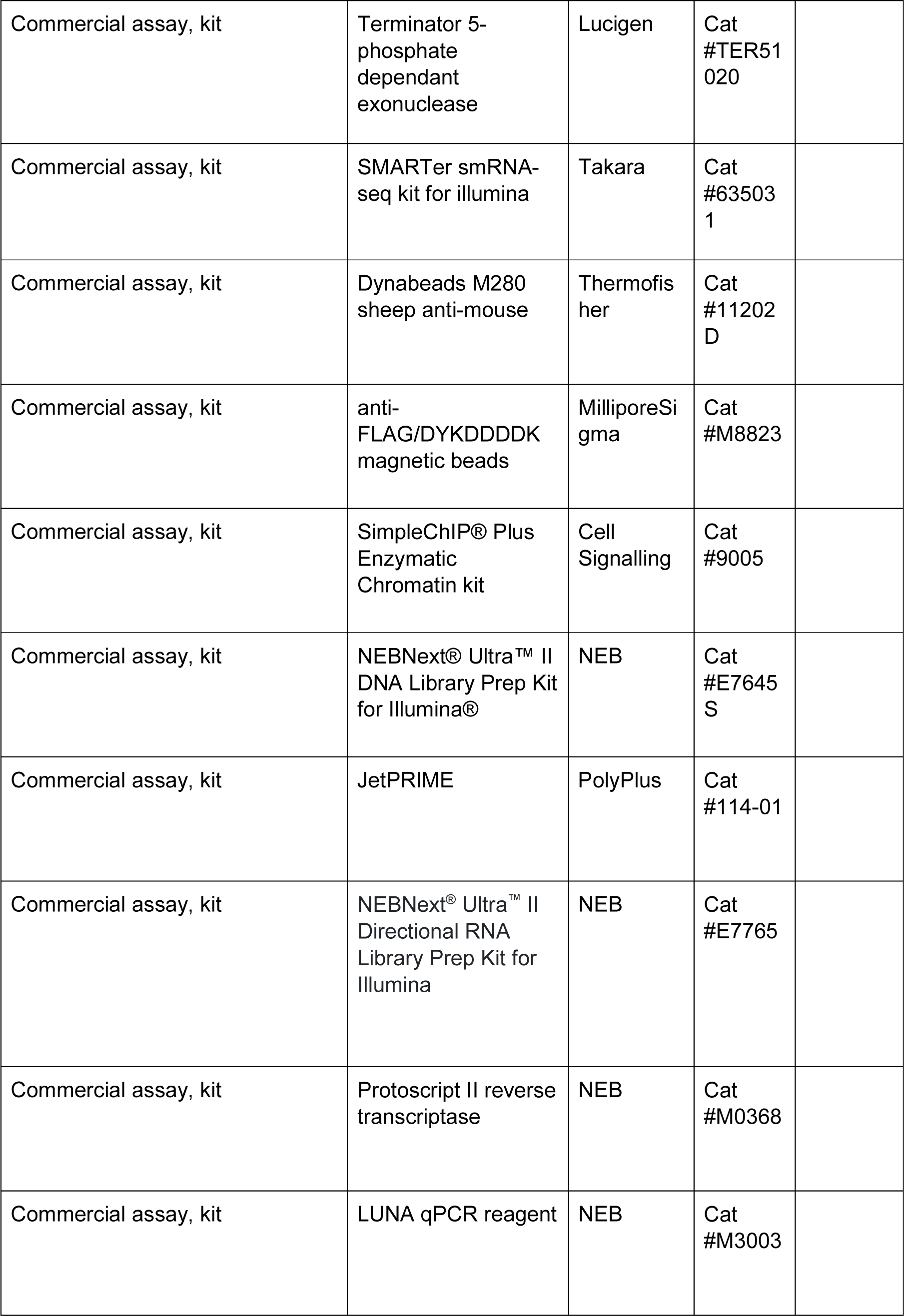
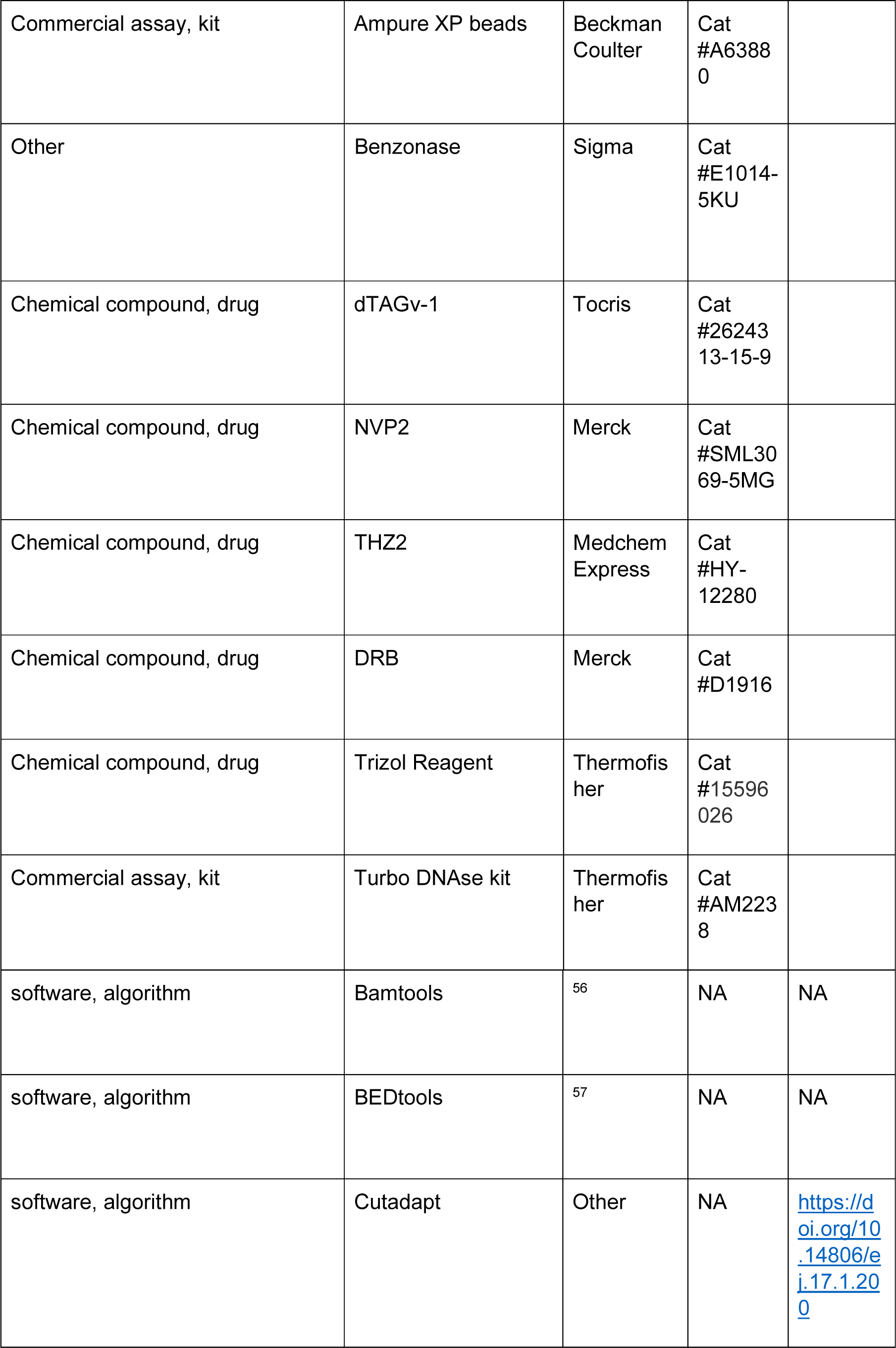
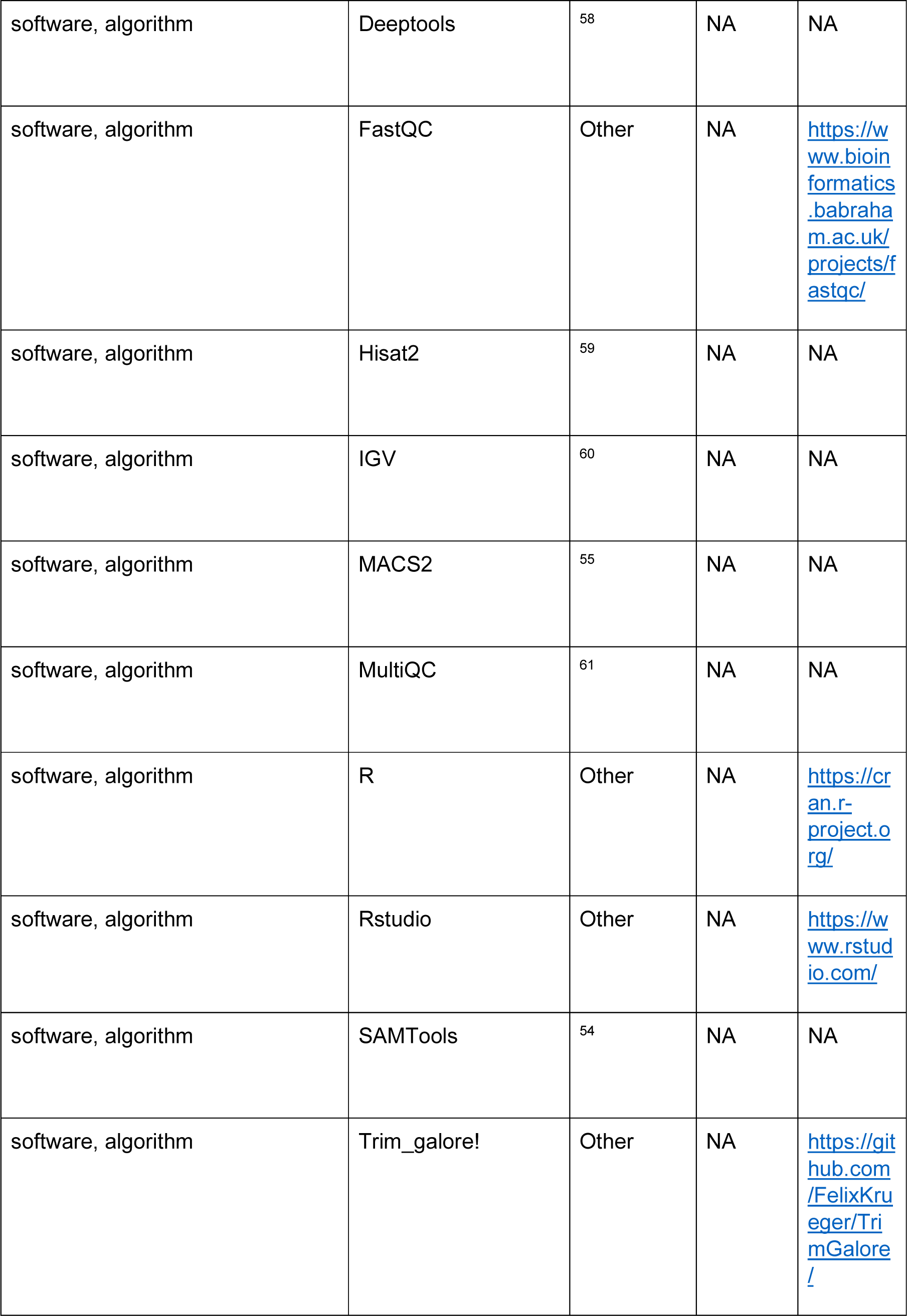
Key Resources Table.

| Key Resources Table |  |  |  |  |
| --- | --- | --- | --- | --- |
| Reagent type (species) or resource | Designation | Source or reference | Identifiers | Additional information |
| cell line (*D. melanogaster*) | S2 | This paper | FLYB:F Btc0000181; RRID:CVCL_Z992 | Cell line maintained in N. Perrimon lab; FlyBase symbol: |
|  |  |  |  | S2-DRSC. |
| Cell line (H. Sapiens) | HCT116 | In house | This paper |  |
| Cell line (H. Sapiens) | *INTS11-dTAG* | In house | This paper |  |
| Cell line (H. Sapiens) | *dTAG-RBBP6* | In house | This paper |  |
| Antibody | INTS11 | Abbexa | abx234340 | WB: 1:500 |
| Antibody | INTS10 | Proteintech | Cat# 15271-1-AP<br>RRID:AB_2127260 | WB: 1:3000 |
| Antibody | INTS8 | Proteintech | Cat# 18802-1-AP<br>RRID:AB_10597250 | WB: 1:1000 |
| Antibody | INTS9 | Proteintech | Cat# 11657-1-AP<br>RRID:AB_2127514 | WB: 1:1000 |
| Antibody | INTS1 | Bethyl | Cat# A300-361A-T<br>RRID:A | WB: 1:500 |
|  |  |  | B_2632<br>121<br>(Now discontinued) |  |
| Antibody | HA | Cell Signalling | Cat# 3724<br>RRID:A<br>B_1549<br>585 | WB:<br>1:2000 |
| Antibody | RNAPII Ser2p | Active Motif | Cat# 61084<br>RRID:A<br>B_2687<br>450 | WB:<br>1:2000 |
| Antibody | RNAPII Ser5p | Millipore | Cat# 04-1572-I<br>RRID:A<br>B_2801<br>296 | WB:<br>1:2000 |
| Antibody | Total RNAPII | Hiroshi Kimura Lab | N/A | WB:<br>1:10,000<br>IP: 5-10ug<br>(Ab:Beads) |
| Antibody | SPT5 | Santa Cruz Biotech | Cat# sc-136075<br>RRID:A<br>B_2286<br>723 | WB:<br>1:1000 |
| Antibody | Flag | Proteintech | Cat# 20543-1-AP<br>RRID:A<br>B_1123<br>2216 | WB:<br>1:5000 |
| Antibody | anti-Rabbit | Proteintech | Cat# SA00001-2<br>RRID:AB_2722564 | WB: 1:2000 |
| Antibody | anti-Mouse | Proteintech | Cat# SA00001-1-A<br>RRID:AB_2890995 | WB: 1:2000 |
| Antibody | anti-Rat | ThermoFisher | Cat #A18865<br>RRID:AB_2535642 | WB: 1:2000 |
| Recombinant DNA reagent | px300 | Addgene | RRID #: Addgene_42230 |  |
| Recombinant DNA reagent | pCRIS-PITChv2-BSD-dTAG (BRD4) | Addgene | RRID: Addgene_91792 |  |
| Recombinant DNA reagent | pCRIS-PITChv2-PURO-dTAG (BRD4) | Addgene | RRID: Addgene_91793 |  |
| Transfected construct | TAT | 52 |  |  |
| Transfected construct | INTS11 WT and E203Q | Twist | This paper | Supplemental Table 1 |
| Transfected construct | HIV reporter | In house | This paper |  |
| Commercial assay, kit | Terminator 5-phosphate dependant exonuclease | Lucigen | Cat #TER51020 |  |
| Commercial assay, kit | SMARTer smRNA-seq kit for illumina | Takara | Cat #635031 |  |
| Commercial assay, kit | Dynabeads M280 sheep anti-mouse | Thermofisher | Cat #11202D |  |
| Commercial assay, kit | anti-FLAG/DYKDDDDK magnetic beads | MilliporeSigma | Cat #M8823 |  |
| Commercial assay, kit | SimpleChIP® Plus Enzymatic Chromatin kit | Cell Signalling | Cat #9005 |  |
| Commercial assay, kit | NEBNext® Ultra™ II DNA Library Prep Kit for Illumina® | NEB | Cat #E7645S |  |
| Commercial assay, kit | JetPRIME | PolyPlus | Cat #114-01 |  |
| Commercial assay, kit | NEBNext® Ultra™ II Directional RNA Library Prep Kit for Illumina | NEB | Cat #E7765 |  |
| Commercial assay, kit | Protoscript II reverse transcriptase | NEB | Cat #M0368 |  |
| Commercial assay, kit | LUNA qPCR reagent | NEB | Cat #M3003 |  |
| Commercial assay, kit | Ampure XP beads | Beckman Coulter | Cat #A63880 |  |
| Other | Benzonase | Sigma | Cat #E1014-5KU |  |
| Chemical compound, drug | dTAGv-1 | Tocris | Cat #2624313-15-9 |  |
| Chemical compound, drug | NVP2 | Merck | Cat #SML3069-5MG |  |
| Chemical compound, drug | THZ2 | Medchem Express | Cat #HY-12280 |  |
| Chemical compound, drug | DRB | Merck | Cat #D1916 |  |
| Chemical compound, drug | Trizol Reagent | Thermofisher | Cat #15596026 |  |
| Commercial assay, kit | Turbo DNase kit | Thermofisher | Cat #AM2238 |  |
| software, algorithm | Bamtools | <sup>56</sup> | NA | NA |
| software, algorithm | BEDtools | <sup>57</sup> | NA | NA |
| software, algorithm | Cutadapt | Other | NA | https://doi.org/10.14806/ej.17.1.200 |
| software, algorithm | Deeptools | 58 | NA | NA |
| software, algorithm | FastQC | Other | NA | https://www.bioinformatics.babraham.ac.uk/projects/fastqc/ |
| software, algorithm | Hisat2 | 59 | NA | NA |
| software, algorithm | IGV | 60 | NA | NA |
| software, algorithm | MACS2 | 55 | NA | NA |
| software, algorithm | MultiQC | 61 | NA | NA |
| software, algorithm | R | Other | NA | https://cran.r-project.org/ |
| software, algorithm | Rstudio | Other | NA | https://www.rstudio.com/ |
| software, algorithm | SAMTools | 54 | NA | NA |
| software, algorithm | Trim_galore! | Other | NA | https://github.com/FelixKrueger/TrimGalore/ |

---

**SUPPLEMENTAL FIGURE 1, related to figure 1.**
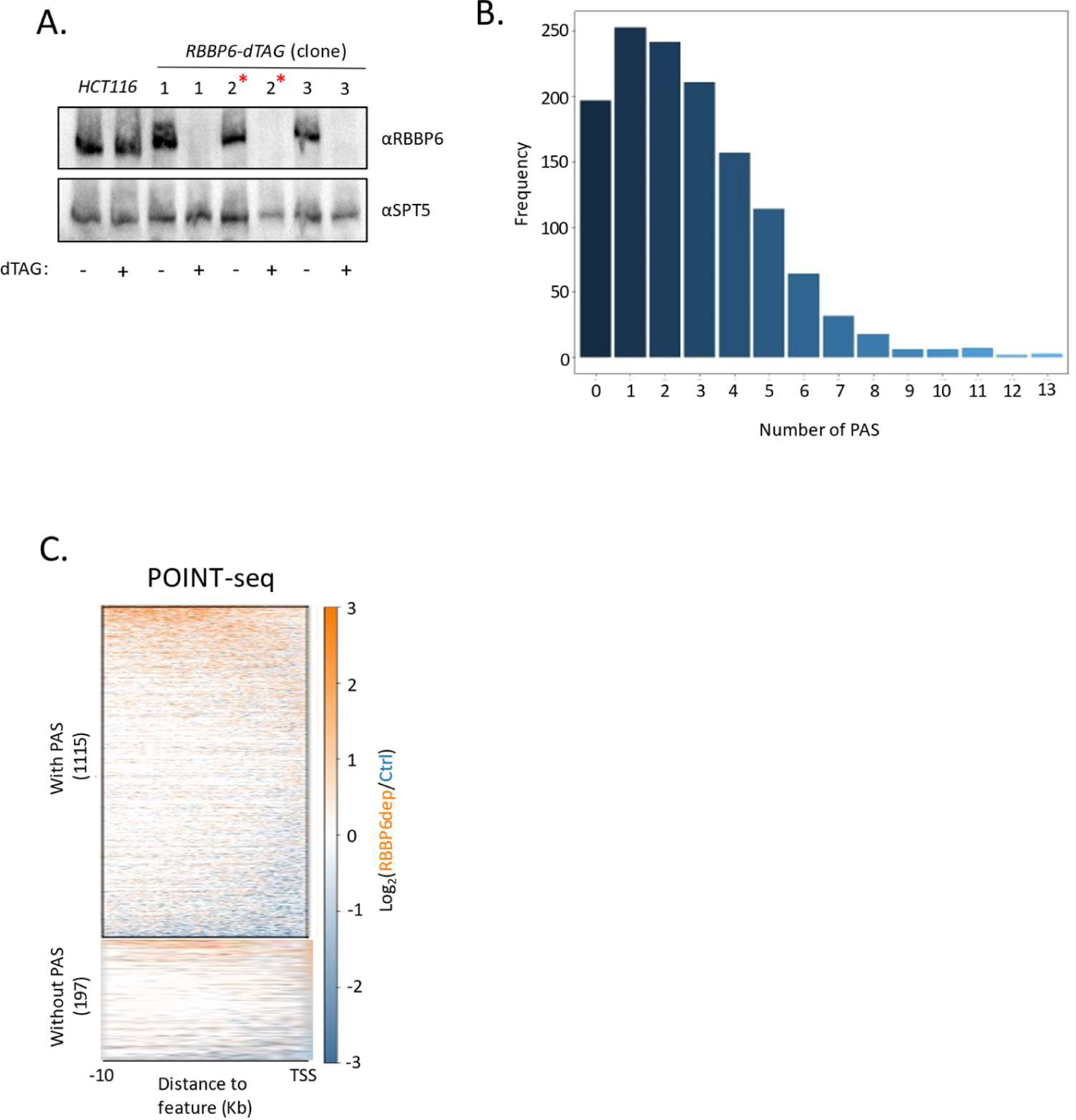
**A**. Western blot showing three separate *dTAG-RBBP6* cell clones. In each case, homozygous tagging is demonstrated by the size shift versus endogenous RBBP6 (HCT116 lanes). Tagged RBBP6 is completely depleted by 2hr treatment with dTAGv-1 whereas endogenous RBBP6 is unaffected. SPT5 serves as a loading control. Clone number 2 (red asterisk) was selected for the experiments in Figure 1. **B.** Graph plotting the number of AAUAAA sequences in antisense transcripts derived from a 3kb window upstream of TSSs. Y-axis is the number of transcripts, and the x-axis shows the AAUAAA count per transcript. **C.** Heatmap of control or RBBP6-depleted POINT-seq showing the RBBP6 effect on antisense transcripts without an AAUAAA or those that contain at least one AAUAAA. Most are unaffected by RBBP6 loss. The y-axis is RPKM.

**SUPPLEMENTAL FIGURE 2, related to figure 2.**
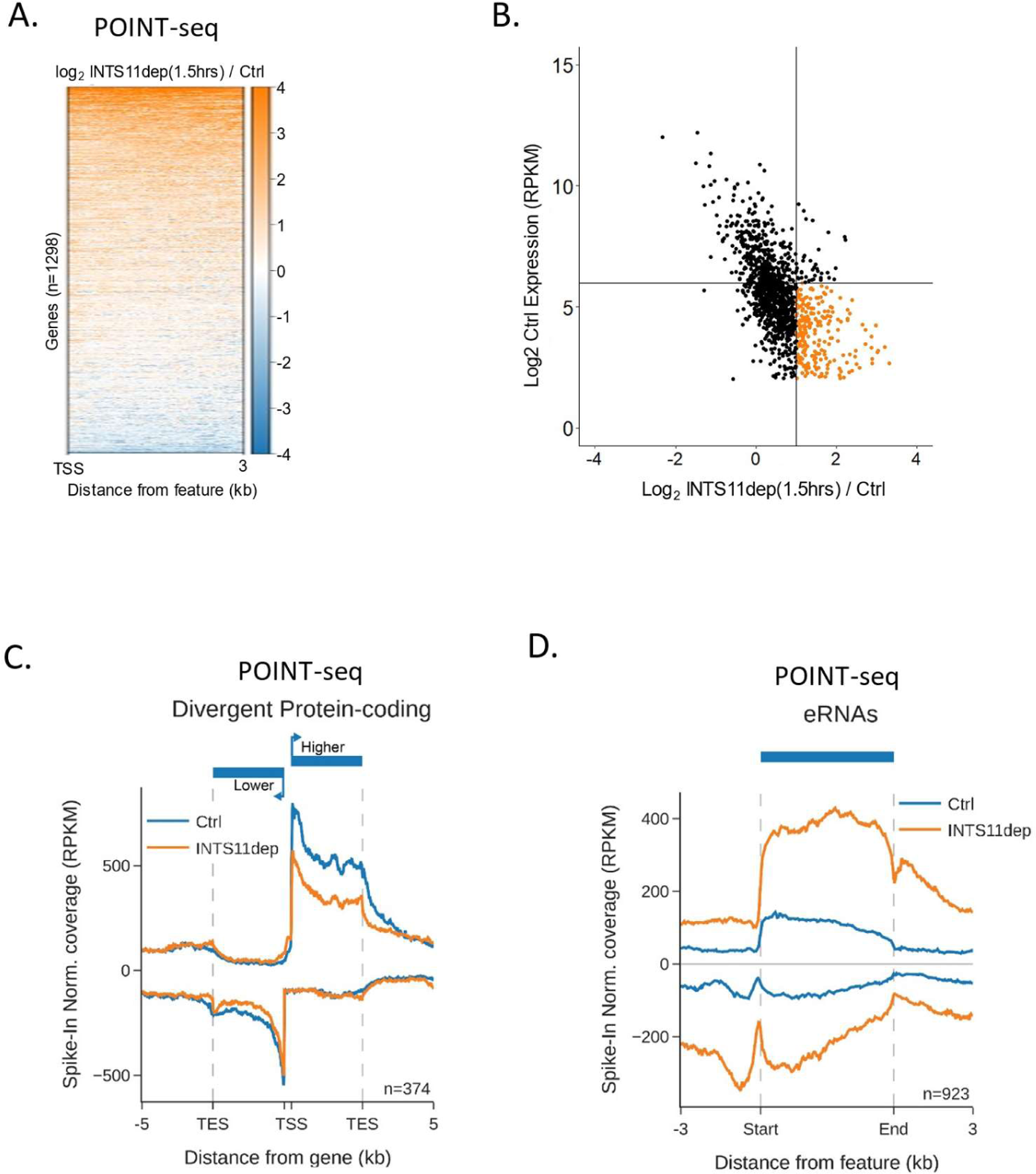
**A.** Heatmap analysis of the 1316 gene set used in Figure 1C but focused on promoter-proximal transcription of the sense direction. Note the change in scale (Log2FC - INTS11/+INTS11) to reflect the smaller effect sizes vs. those for antisense transcription from the same gene set. **B.** Comparison of INTS11-dependent changes in promoter-proximal protein-coding transcription with the expression level of each gene using the 1316 gene set as Figure 1C. X-axis shows Log2FC in levels (-INTS11/+INTS11) and y axis shows expression level. Note that genes with the largest increase in promoter-proximal signal following INTS11 loss are low expressed (coloured orange). Data derives from POINT-seq following treatment or not with dTAGv-1 (1.5hr). **C.** Metaplot of POINT-seq data from *INTS11-dTAG* cells treated or not (1.5hr) with dTAGv-1. These are protein-coding genes arranged head-to-head. Signals above and below the x-axis are sense and antisense reads, respectively. Y-axis shows RPKM following spike-in normalisation. **D.** Metaplot of POINT-seq data from *INTS11-dTAG* cells treated or not (1.5hr) with dTAGv-1. This shows RNAs derived from enhancer clusters, which generally initiate transcription bidirectionally. Signals above and below the x-axis are sense and antisense reads, respectively. Y-axis shows RPKM following spike-in normalisation.

**SUPPLEMENTAL FIGURE 3, related to figure 3.**
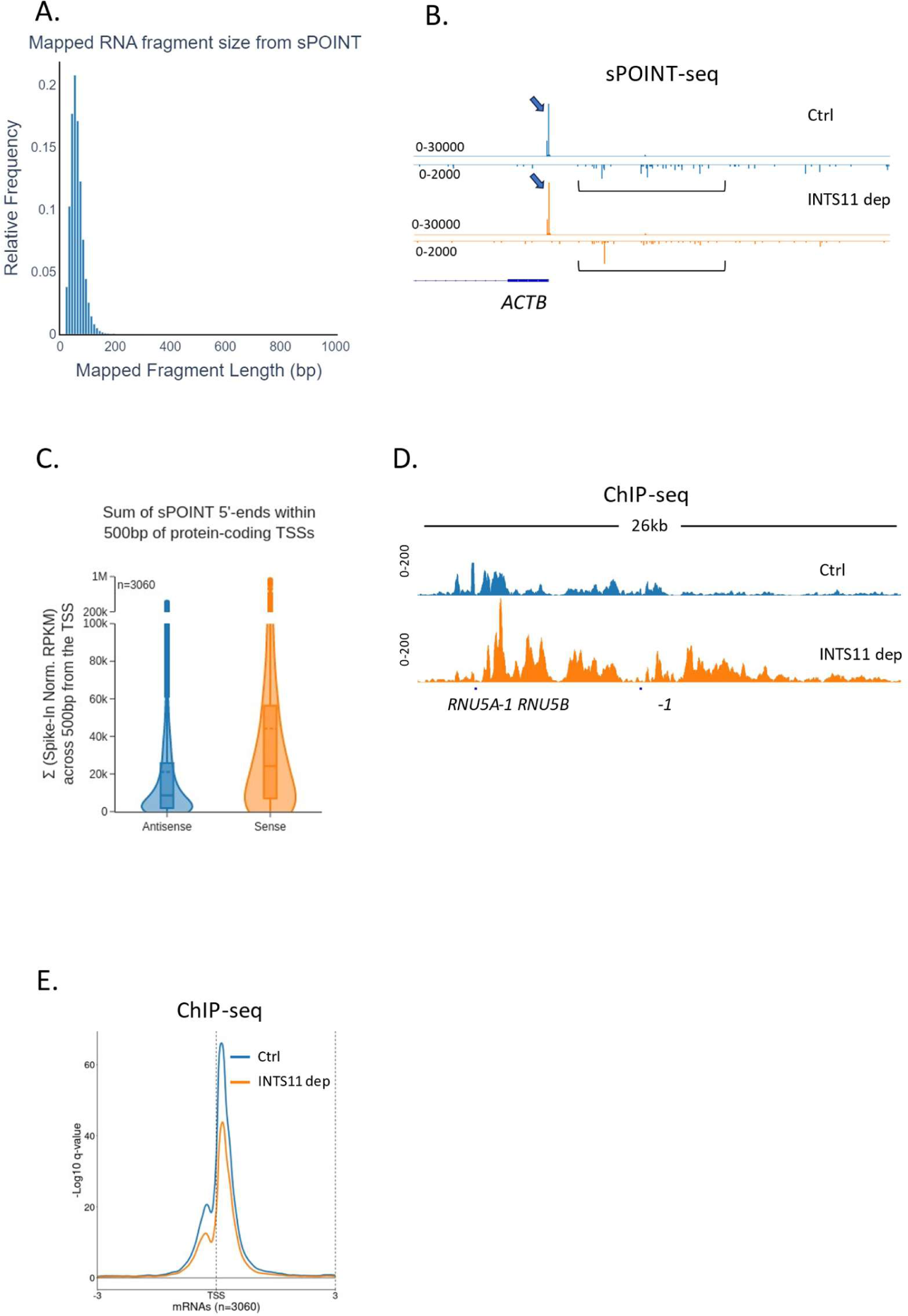
**A.** Plot showing the size distribution of fragments mapped by sPOINT-seq. This demonstrates its selectivity toward transcripts <150nts. **B**. Genome browser track of *ACTB* promoter region in sPOINT-seq from *INTS11-dTAG* cells treated or not with dTAGv-1 (1.5 hr). This showcases the focused sense TSS (black arrows) and the dispersed antisense reads (brackets). Note the different y-axis scales (RPKM) for sense vs. antisense. **C.** Violin plot of the sum sPOINT 5’end spike-in normalised RPKM signal across a 500bp window from the protein-coding TSSs shown in Figure 3D (n=3060). These samples are untreated with dTAG to quantitate the normal levels of sense vs. antisense initiation/pausing in a window -/+ 500bp from the TSS in respective directions for antisense and sense. This shows more initiation/pausing in sense vs. antisense directions. **D**. Genome track of *RNU5A-1* and *RNU5B-1* in RNAPII ChIP-seq performed on *INTS11-dTAG* cells treated or not with dTAGv-1 (2hr). INTS11 depletion causes RNAPII build-up beyond both genes indicating a transcription termination defect. The Y-axis scale is RPKM. **E**. Metaplot of RNAPII occupancy of protein-coding TSS regions (the same gene set employed for the sPOINT analysis in Figure 3) derived from RNAPII ChIP-seq performed on *INTS11-dTAG* cells treated or not with dTAGv-1 (2hr). The Y-axis scale is log10 qvalue of peak pileup values normalised to spike in control.

**SUPPLEMENTAL FIGURE 4, related to figure 4.**
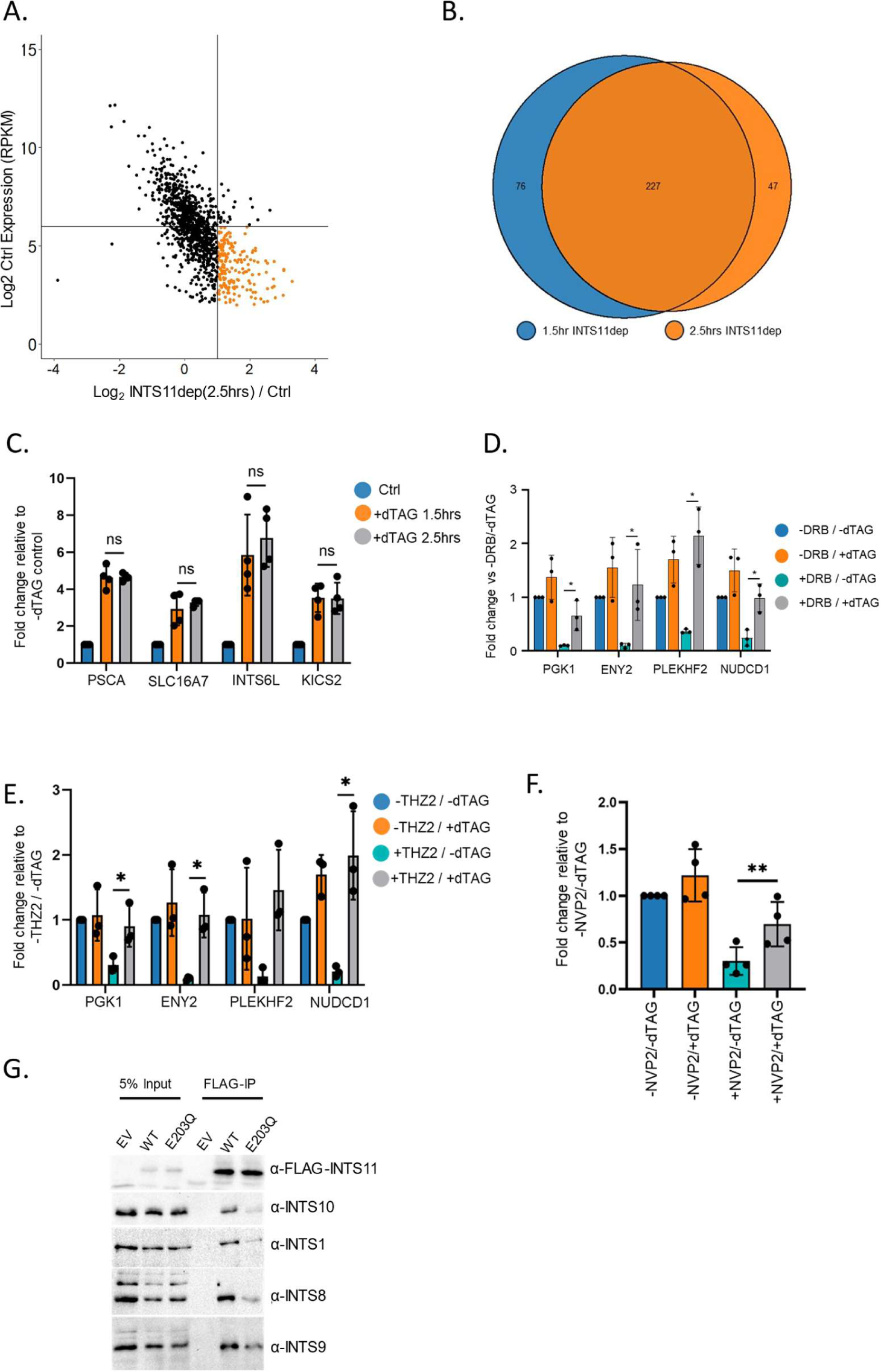
**A**. Comparison of INTS11-dependent changes in promoter-proximal protein-coding transcription with the expression level of each gene. The gene set is equivalent to the analogous panel in Supplemental Figure 2B. The x-axis shows Log2FC in levels (-INTS11/+INTS11) and the y-axis shows expression level. Note that genes with the largest increase in promoter-proximal signal following INTS11 loss are low expressed (coloured orange). Data derives from POINT-seq following treatment or not with dTAGv-1 (2.5hr). **B.** Venn diagram showing the number of protein-coding genes where promoter-proximal POINT-seq signal increases by ≥Log2FC of 1 after INTS11 loss after 1.5h or 2.5h dTAGv-1 treatment. A strong overlap is seen indicating reproducible effects at the two time points. **C.** qRT-PCR analysis of PSCA, SLC16A7, INTS6L, and KICS2 pre-mRNAs in *INTS11-dTAG* cells treated or not with dTAGv-1 for 1.5hr or 2.5hr. To enrich nascent transcripts, primers detect intronic RNA. Quantitation shows fold change versus spliced actin relative to untreated samples. Error bars show standard deviation. n=4. n.s denotes not significant. **D.** qRT-PCR analysis of PGK1, ENY2, PLEKHF2, and NUDCD1 pre-mRNAs in *INTS11-dTAG* cells treated or not with dTAGv-1 and at the same time exposed or not to DRB (all 2.5hr). To enrich nascent transcripts, primers detect intronic RNA. Quantitation shows fold change versus spliced actin relative to samples untreated with dTAGv-1 or DRB. DRB treatment substantially reduces signal, which is restored when INTS11 is co-depleted. Error bars show standard deviation. n=3, * denotes p≤0.05. **E.** As for C but following treatment with the CDK7 inhibitor, THZ1, and/or depletion of INTS11 with dTAGv-1 (all treatments 2.5hr). Error bars show standard deviation. n=3, * denotes p≤0.05. **F.** qRT-PCR analysis of chromatin-associated (nascent) RNA isolated from *INTS11-dTAG* cells transfected with the HIV reporter construct together with TAT then treated or not with INTS11 and/or NVP-2. Quantitation shows signals relative to those obtained in untreated cells after normalising to MALAT1 RNA. n=4. Error bars show standard deviation. **denotes p≤0.01. **G.** Flag immunoprecipitation performed in *INTS11-dTAG* cells transfected with flag-tagged wild-type or E203Q INTS11. The gel shows input and co-precipitated samples probed for components of different Integrator modules (tail module – INTS10; backbone module – INTS1; phosphatase module – INTS8 and cleavage module – INTS9). These components are poorly associated with E203Q INTS11 vs. wild type.

## REFERENCES

1. Preker, P., Nielsen, J., Kammler, S., Lykke-Andersen, S., Christensen, M.S., Mapendano, C.K., Schierup, M.H., and Jensen, T.H. (2008). RNA exosome depletion reveals transcription upstream of active human promoters. Science 322, 1851–1854. 1164096 [pii] 10.1126/science.1164096.

2. Wyers, F., Rougemaille, M., Badis, G., Rousselle, J.C., Dufour, M.E., Boulay, J., Regnault, B., Devaux, F., Namane, A., Seraphin, B., et al. (2005). Cryptic pol II transcripts are degraded by a nuclear quality control pathway involving a new poly(A) polymerase. Cell 121, 725–737. 10.1016/j.cell.2005.04.030.

3. Thieffry, A., Vigh, M.L., Bornholdt, J., Ivanov, M., Brodersen, P., and Sandelin, A. (2020). Characterization of Arabidopsis thaliana Promoter Bidirectionality and Antisense RNAs by Inactivation of Nuclear RNA Decay Pathways. Plant Cell 32, 1845–1867. 10.1105/tpc.19.00815.

4. Jin, Y., Eser, U., Struhl, K., and Churchman, L.S. (2017). The Ground State and Evolution of Promoter Region Directionality. Cell 170, 889–898 e810. 10.1016/j.cell.2017.07.006.

5. Kaida, D., Berg, M.G., Younis, I., Kasim, M., Singh, L.N., Wan, L., and Dreyfuss, G. (2010). U1 snRNP protects pre-mRNAs from premature cleavage and polyadenylation. Nature 468, 664–668. nature09479 [pii] 10.1038/nature09479.

6. Mimoso, C.A., and Adelman, K. (2023). U1 snRNP increases RNA Pol II elongation rate to enable synthesis of long genes. Molecular cell. 10.1016/j.molcel.2023.03.002.

7. Chiu, A.C., Suzuki, H.I., Wu, X., Mahat, D.B., Kriz, A.J., and Sharp, P.A. (2018). Transcriptional Pause Sites Delineate Stable Nucleosome-Associated Premature Polyadenylation Suppressed by U1 snRNP. Molecular cell 69, 648–663 e647. 10.1016/j.molcel.2018.01.006.

8. So, B.R., Di, C., Cai, Z., Venters, C.C., Guo, J., Oh, J.M., Arai, C., and Dreyfuss, G. (2019). A Complex of U1 snRNP with Cleavage and Polyadenylation Factors Controls Telescripting, Regulating mRNA Transcription in Human Cells. Molecular cell 76, 590–599 e594. 10.1016/j.molcel.2019.08.007.

9. Estell, C., Davidson, L., Eaton, J.D., Kimura, H., Gold, V.A.M., and West, S. (2023). A restrictor complex of ZC3H4, WDR82, and ARS2 integrates with PNUTS to control unproductive transcription. Molecular cell 83, 2222-2239 e2225. 10.1016/j.molcel.2023.05.029.

10. Almada, A.E., Wu, X., Kriz, A.J., Burge, C.B., and Sharp, P.A. (2013). Promoter directionality is controlled by U1 snRNP and polyadenylation signals. Nature 499, 360–363. 10.1038/nature12349.

11. Wagner, E.J., Tong, L., and Adelman, K. (2023). Integrator is a global promoter-proximal termination complex. Molecular cell 83, 416–427. 10.1016/j.molcel.2022.11.012.

12. Lykke-Andersen, S., Zumer, K., Molska, E.S., Rouviere, J.O., Wu, G., Demel, C., Schwalb, B., Schmid, M., Cramer, P., and Jensen, T.H. (2021). Integrator is a genome-wide attenuator of non-productive transcription. Molecular cell 81, 514–529 e516. 10.1016/j.molcel.2020.12.014.

13. Beckedorff, F., Blumenthal, E., daSilva, L.F., Aoi, Y., Cingaram, P.R., Yue, J., Zhang, A., Dokaneheifard, S., Valencia, M.G., Gaidosh, G., et al. (2020). The Human Integrator Complex Facilitates Transcriptional Elongation by Endonucleolytic Cleavage of Nascent Transcripts. Cell reports 32, 107917. 10.1016/j.celrep.2020.107917.

14. Tatomer, D.C., Elrod, N.D., Liang, D., Xiao, M.S., Jiang, J.Z., Jonathan, M., Huang, K.L., Wagner, E.J., Cherry, S., and Wilusz, J.E. (2019). The Integrator complex cleaves nascent mRNAs to attenuate transcription. Genes & development 33, 1525–1538. 10.1101/gad.330167.119.

15. Elrod, N.D., Henriques, T., Huang, K.L., Tatomer, D.C., Wilusz, J.E., Wagner, E.J., and Adelman, K. (2019). The Integrator Complex Attenuates Promoter-Proximal Transcription at Protein-Coding Genes. Molecular cell 76, 738–752 e737. 10.1016/j.molcel.2019.10.034.

16. Hu, S., Peng, L., Song, A., Ji, Y.X., Cheng, J., Wang, M., and Chen, F.X. (2023). INTAC endonuclease and phosphatase modules differentially regulate transcription by RNA polymerase II. Molecular cell 83, 1588–1604 e1585. 10.1016/j.molcel.2023.03.022.

17. Stein, C.B., Field, A.R., Mimoso, C.A., Zhao, C., Huang, K.L., Wagner, E.J., and Adelman, K. (2022). Integrator endonuclease drives promoter-proximal termination at all RNA polymerase II-transcribed loci. Molecular cell 82, 4232–4245 e4211. 10.1016/j.molcel.2022.10.004.

18. Vervoort, S.J., Welsh, S.A., Devlin, J.R., Barbieri, E., Knight, D.A., Offley, S., Bjelosevic, S., Costacurta, M., Todorovski, I., Kearney, C.J., et al. (2021). The PP2A-Integrator-CDK9 axis fine-tunes transcription and can be targeted therapeutically in cancer. Cell 184, 3143–3162 e3132. 10.1016/j.cell.2021.04.022.

19. Gockert, M., Schmid, M., Jakobsen, L., Jens, M., Andersen, J.S., and Jensen, T.H. (2022). Rapid factor depletion highlights intricacies of nucleoplasmic RNA degradation. Nucleic acids research 50, 1583–1600. 10.1093/nar/gkac001.

20. Schmidt, M., Kluge, F., Sandmeir, F., Kuhn, U., Schafer, P., Tuting, C., Ihling, C., Conti, E., and Wahle, E. (2022). Reconstitution of 3’ end processing of mammalian pre-mRNA reveals a central role of RBBP6. Genes & development 36, 195–209. 10.1101/gad.349217.121.

21. Boreikaite, V., Elliott, T.S., Chin, J.W., and Passmore, L.A. (2022). RBBP6 activates the pre-mRNA 3’ end processing machinery in humans. Genes & development 36, 210–224. 10.1101/gad.349223.121.

22. Sousa-Luis, R., Dujardin, G., Zukher, I., Kimura, H., Weldon, C., Carmo-Fonseca, M., Proudfoot, N.J., and Nojima, T. (2021). POINT technology illuminates the processing of polymerase-associated intact nascent transcripts. Molecular cell 81, 1935–1950 e1936. 10.1016/j.molcel.2021.02.034.

23. Baillat, D., Hakimi, M.A., Naar, A.M., Shilatifard, A., Cooch, N., and Shiekhattar, R. (2005). Integrator, a multiprotein mediator of small nuclear RNA processing, associates with the C-terminal repeat of RNA polymerase II. Cell 123, 265–276. 10.1016/j.cell.2005.08.019.

24. Nabet, B., Roberts, J.M., Buckley, D.L., Paulk, J., Dastjerdi, S., Yang, A., Leggett, A.L., Erb, M.A., Lawlor, M.A., Souza, A., et al. (2018). The dTAG system for immediate and target-specific protein degradation. Nature chemical biology 14, 431–441. 10.1038/s41589-018-0021-8.

25. Zheng, H., Qi, Y., Hu, S., Cao, X., Xu, C., Yin, Z., Chen, X., Li, Y., Liu, W., Li, J., et al. (2020). Identification of Integrator-PP2A complex (INTAC), an RNA polymerase II phosphatase. Science 370. 10.1126/science.abb5872.

26. Ramamurthy, L., Ingledue, T.C., Pilch, D.R., Kay, B.K., and Marzluff, W.F. (1996). Increasing the distance between the snRNA promoter and the 3’ box decreases the efficiency of snRNA 3’-end formation. Nucleic acids research 24, 4525–4534. 10.1093/nar/24.22.4525.

27. Fujinaga, K., Huang, F., and Peterlin, B.M. (2023). P-TEFb: The master regulator of transcription elongation. Molecular cell 83, 393–403. 10.1016/j.molcel.2022.12.006.

28. Olson, C.M., Jiang, B., Erb, M.A., Liang, Y., Doctor, Z.M., Zhang, Z., Zhang, T., Kwiatkowski, N., Boukhali, M., Green, J.L., et al. (2018). Pharmacological perturbation of CDK9 using selective CDK9 inhibition or degradation. Nature chemical biology 14, 163–170. 10.1038/nchembio.2538.

29. Schuller, R., Forne, I., Straub, T., Schreieck, A., Texier, Y., Shah, N., Decker, T.M., Cramer, P., Imhof, A., and Eick, D. (2016). Heptad-Specific Phosphorylation of RNA Polymerase II CTD. Molecular cell 61, 305–314. 10.1016/j.molcel.2015.12.003.

30. Suh, H., Ficarro, S.B., Kang, U.B., Chun, Y., Marto, J.A., and Buratowski, S. (2016). Direct Analysis of Phosphorylation Sites on the Rpb1 C-Terminal Domain of RNA Polymerase II. Molecular cell 61, 297–304. 10.1016/j.molcel.2015.12.021.

31. Larochelle, S., Amat, R., Glover-Cutter, K., Sanso, M., Zhang, C., Allen, J.J., Shokat, K.M., Bentley, D.L., and Fisher, R.P. (2012). Cyclin-dependent kinase control of the initiation-to-elongation switch of RNA polymerase II. Nat Struct Mol Biol 19, 1108–1115. 10.1038/nsmb.2399.

32. Wei, P., Garber, M.E., Fang, S.M., Fischer, W.H., and Jones, K.A. (1998). A novel CDK9-associated C-type cyclin interacts directly with HIV-1 Tat and mediates its high-affinity, loop-specific binding to TAR RNA. Cell 92, 451–462. 10.1016/s0092-8674(00)80939-3.

33. Stadelmayer, B., Micas, G., Gamot, A., Martin, P., Malirat, N., Koval, S., Raffel, R., Sobhian, B., Severac, D., Rialle, S., et al. (2014). Integrator complex regulates NELF-mediated RNA polymerase II pause/release and processivity at coding genes. Nat Commun 5, 5531. 10.1038/ncomms6531.

34. Rengachari, S., Schilbach, S., Kaliyappan, T., Gouge, J., Zumer, K., Schwarz, J., Urlaub, H., Dienemann, C., Vannini, A., and Cramer, P. (2022). Structural basis of SNAPc-dependent snRNA transcription initiation by RNA polymerase II. Nat Struct Mol Biol 29, 1159–1169. 10.1038/s41594-022-00857-w.

35. Kolev, N.G., Yario, T.A., Benson, E., and Steitz, J.A. (2008). Conserved motifs in both CPSF73 and CPSF100 are required to assemble the active endonuclease for histone mRNA 3’-end maturation. EMBO Rep 9, 1013–1018. 10.1038/embor.2008.146.

36. Fianu, I., Chen, Y., Dienemann, C., Dybkov, O., Linden, A., Urlaub, H., and Cramer, P. (2021). Structural basis of Integrator-mediated transcription regulation. Science 374, 883–887. 10.1126/science.abk0154.

37. Yamada, T., Yamaguchi, Y., Inukai, N., Okamoto, S., Mura, T., and Handa, H. (2006). P-TEFb-mediated phosphorylation of hSpt5 C-terminal repeats is critical for processive transcription elongation. Molecular cell 21, 227–237. S1097-2765(05)01812-5 [pii] 10.1016/j.molcel.2005.11.024.

38. Cortazar, M.A., Sheridan, R.M., Erickson, B., Fong, N., Glover-Cutter, K., Brannan, K., and Bentley, D.L. (2019). Control of RNA Pol II Speed by PNUTS-PP1 and Spt5 Dephosphorylation Facilitates Termination by a “Sitting Duck Torpedo” Mechanism. Molecular cell 76, 896–908 e894. 10.1016/j.molcel.2019.09.031.

39. Parua, P.K., Booth, G.T., Sanso, M., Benjamin, B., Tanny, J.C., Lis, J.T., and Fisher, R.P. (2018). A Cdk9-PP1 switch regulates the elongation-termination transition of RNA polymerase II. Nature 558, 460–464. 10.1038/s41586-018-0214-z.

40. Fong, N., Sheridan, R.M., Ramachandran, S., and Bentley, D.L. (2022). The pausing zone and control of RNA polymerase II elongation by Spt5: Implications for the pause-release model. Molecular cell 82, 3632–3645 e3634. 10.1016/j.molcel.2022.09.001.

41. Flynn, R.A., Almada, A.E., Zamudio, J.R., and Sharp, P.A. (2011). Antisense RNA polymerase II divergent transcripts are P-TEFb dependent and substrates for the RNA exosome. Proceedings of the National Academy of Sciences of the United States of America 108, 10460–10465. 10.1073/pnas.1106630108.

42. Wu, G., Schmid, M., Rib, L., Polak, P., Meola, N., Sandelin, A., and Jensen, T.H. (2020). A Two-Layered Targeting Mechanism Underlies Nuclear RNA Sorting by the Human Exosome. Cell reports 30, 2387–2401 e2385. 10.1016/j.celrep.2020.01.068.

43. Meola, N., Domanski, M., Karadoulama, E., Chen, Y., Gentil, C., Pultz, D., Vitting-Seerup, K., Lykke-Andersen, S., Andersen, J.S., Sandelin, A., and Jensen, T.H. (2016). Identification of a Nuclear Exosome Decay Pathway for Processed Transcripts. Molecular cell 64, 520–533. 10.1016/j.molcel.2016.09.025.

44. Estell, C., Davidson, L., Steketee, P.C., Monier, A., and West, S. (2021). ZC3H4 restricts non-coding transcription in human cells. Elife 10. 10.7554/eLife.67305.

45. Austenaa, L.M.I., Piccolo, V., Russo, M., Prosperini, E., Polletti, S., Polizzese, D., Ghisletti, S., Barozzi, I., Diaferia, G.R., and Natoli, G. (2021). A first exon termination checkpoint preferentially suppresses extragenic transcription. Nat Struct Mol Biol 28, 337–346. 10.1038/s41594-021-00572-y.

46. Vlaming, H., Mimoso, C.A., Field, A.R., Martin, B.J.E., and Adelman, K. (2022). Screening thousands of transcribed coding and non-coding regions reveals sequence determinants of RNA polymerase II elongation potential. Nat Struct Mol Biol. 10.1038/s41594-022-00785-9.

47. Muniz, L., Davidson, L., and West, S. (2015). Poly(A) Polymerase and the Nuclear Poly(A) Binding Protein, PABPN1, Coordinate the Splicing and Degradation of a Subset of Human Pre-mRNAs. Molecular and cellular biology 35, 2218-2230. 10.1128/MCB.00123-15.

48. Dye, M.J., and Proudfoot, N.J. (2001). Multiple transcript cleavage precedes polymerase release in termination by RNA polymerase II. Cell 105, 669–681. S0092-8674(01)00372-5 [pii].

49. Davidson, L., Francis, L., Eaton, J.D., and West, S. (2020). Integrator-Dependent and Allosteric/Intrinsic Mechanisms Ensure Efficient Termination of snRNA Transcription. Cell reports 33, 108319. 10.1016/j.celrep.2020.108319.

50. Cong, L., Ran, F.A., Cox, D., Lin, S., Barretto, R., Habib, N., Hsu, P.D., Wu, X., Jiang, W., Marraffini, L.A., and Zhang, F. (2013). Multiplex genome engineering using CRISPR/Cas systems. Science 339, 819–823. science.1231143 [pii] 10.1126/science.1231143.

51. Manna, P.T., Davis, L.J., and Robinson, M.S. (2019). Fast and cloning-free CRISPR/Cas9-mediated genomic editing in mammalian cells. Traffic 20, 974–982. 10.1111/tra.12696.

52. Adams, S.E., Johnson, I.D., Braddock, M., Kingsman, A.J., Kingsman, S.M., and Edwards, R.M. (1988). Synthesis of a gene for the HIV transactivator protein TAT by a novel single stranded approach involving in vivo gap repair. Nucleic acids research 16, 4287–4298.

53. Gregersen, L.H., Mitter, R., and Svejstrup, J.Q. (2022). Elongation factor-specific capture of RNA polymerase II complexes. Cell Rep Methods 2, 100368. 10.1016/j.crmeth.2022.100368.

54. Li, H., Handsaker, B., Wysoker, A., Fennell, T., Ruan, J., Homer, N., Marth, G., Abecasis, G., Durbin, R., and Genome Project Data Processing, S. (2009). The Sequence Alignment/Map format and SAMtools. Bioinformatics 25, 2078-2079. 10.1093/bioinformatics/btp352.

55. Zhang, Y., Liu, T., Meyer, C.A., Eeckhoute, J., Johnson, D.S., Bernstein, B.E., Nusbaum, C., Myers, R.M., Brown, M., Li, W., and Liu, X.S. (2008). Model-based analysis of ChIP-Seq (MACS). Genome Biol 9, R137. 10.1186/gb-2008-9-9-r137.

56. Barnett, D.W., Garrison, E.K., Quinlan, A.R., Stromberg, M.P., and Marth, G.T. (2011). BamTools: a C++ API and toolkit for analyzing and managing BAM files. Bioinformatics 27, 1691–1692. 10.1093/bioinformatics/btr174.

57. Quinlan, A.R., and Hall, I.M. (2010). BEDTools: a flexible suite of utilities for comparing genomic features. Bioinformatics 26, 841–842. 10.1093/bioinformatics/btq033.

58. Ramirez, F., Ryan, D.P., Gruning, B., Bhardwaj, V., Kilpert, F., Richter, A.S., Heyne, S., Dundar, F., and Manke, T. (2016). deepTools2: a next generation web server for deep-sequencing data analysis. Nucleic acids research 44, W160–165. 10.1093/nar/gkw257.

59. Kim, D., Langmead, B., and Salzberg, S.L. (2015). HISAT: a fast spliced aligner with low memory requirements. Nat Methods 12, 357–360. 10.1038/nmeth.3317.

60. Robinson, J.T., Thorvaldsdottir, H., Winckler, W., Guttman, M., Lander, E.S., Getz, G., and Mesirov, J.P. (2011). Integrative genomics viewer. Nat Biotechnol 29, 24–26. 10.1038/nbt.1754.

61. Ewels, P., Magnusson, M., Lundin, S., and Kaller, M. (2016). MultiQC: summarize analysis results for multiple tools and samples in a single report. Bioinformatics 32, 3047–3048. 10.1093/bioinformatics/btw354.

